# Variational inference of single cell time series

**DOI:** 10.1101/2024.08.29.610389

**Authors:** Bingxian Xu, Rosemary Braun

## Abstract

Time course single–cell RNA sequencing (scRNA-seq) enables researchers to probe expression dynamics at the resolution of individual cells. However, analyzing this rich data poses several challenges, including deconvolving the contributions of time and cell type, discriminating true dynamics from batch effects, and inferring per–cell dynamics despite cells being destroyed at each time point. We present SNOW (SiNgle cell flOW map), a deep learning algorithm that deconvolves single-cell time series into time-dependent and time-independent components. Using both synthetic and real scRNA-seq data, we show that SNOW constructs biologically meaningful latent spaces, removes batch effects, and generates realistic single-cell time series.

## 1 Background

Gene expression is shaped by intrinsic cellular identities and extrinsic environmental conditions. Today, single-cell RNA sequencing (scRNA-seq) technologies enable us to probe how gene expression changes across cell types under various experimental conditions [1–5], with applications ranging from organ development [6, 7] to cancer progression [8, 9] and more recently to circadian rhythms [10, 11]. To understand the dynamics of these processes, studies have started to directly observe how gene expression profiles change over time via time–coures scRNA-seq profiling [7, 12–14] and a number of methods have been developed to characterize and model scRNA-seq time-series data. For example, Waddington-OT [6] applies unbalanced optimal transport to compute the likelihood of cell state transitions. To gain mechanistic insights, PRESCIENT (Potential eneRgy undErlying Single Cell gradIENTs) [15] constructs a global potential function, *Ψ*(**x**), and uses Δ*Ψ*(**x**) to estimate how gene expression, **x**, changes over time via the Euler scheme **x**(*t* + *δt*) = **x**(*t*) *−* Δ*Ψ*(**x**) *δt*. However, this potential function is constructed on the PCA space, which may not represent the relevant geometry and cannot be mapped back to the original gene expression space after the dimensionality is reduced. If the data lie on a nonlinear/curved manifold, such as in the “Swiss roll” example [16] or when gene expression dynamics have a cyclic component, PCs will fail to articulate this coordinate of variation. To overcome this limitation, scNODE [17] uses a variational autoencoder [18] to construct a lower dimensional space with which to find governing equations that recapitulate the observed dynamics.

All the aforementioned methods are some variant of parameterizing a flow that satisfies the optimal transport constraint. For example, PRESCIENT [15] and scNODE [17] minimize the Wasserstein distance between data sampled from one time point and that sampled from an earlier time point that subsequently evolved according to the flow. However, this treatment only constrains the flow on measured time points. To further constrain the flow, methods such as TIGON [19] and TrajectoryNet [20] solve the dynamic optimal transport problem [21] by regularizing the entire path along which the probability densities evolve. This approach is useful in contexts were temporal variation affects all cells, such as during development where cells move along common paths in a low dimensional space (Figure 1A, top). However, this may not be the best description for systems where cells can act in a highly cell type–specific manner over time (Figure 1A, bottom), which may complicate both cell type annotations and temporal analysis. In this situation, it is desirable to remove the effect of time to facilitate cell type annotation, which is usually achieved by integrating and batch–correcting the time points. Since removing temporal variation also removes biologically meaningful dynamics that one may wish to study, further analyses must use non-integrated data to study the average expression for each cell type over time. This “two step” approach has been used to study cellular responses to perturbations [22] as well as circadian dynamics [10]. However, conclusions drawn from the second stage analyses will depend on the quality of the integration and the resulting clusters / cell type annotation from the first stage. Additionally, data integration remains an active field of research [23–26], and discriminating batch effects from temporal variation is particularly difficult. With the growing interest in time–resolved scRNA-seq profiling, there is a pressing need for methods designed specifically for scRNA time series analysis.

**Figure 1.**
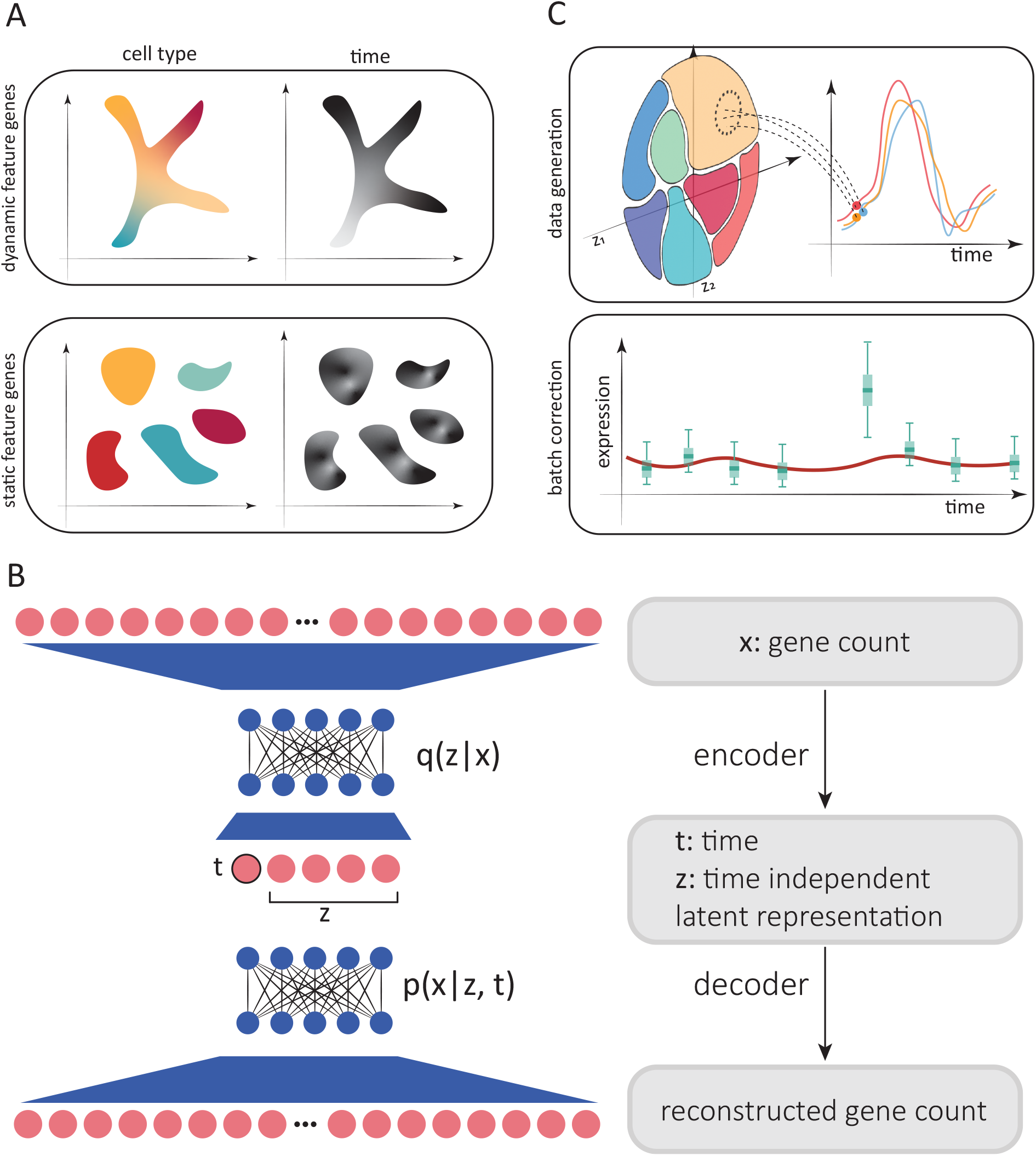
SNOW overview. A: Two possible scenarios of temporal effects in scRNA-seq time–series data. Top: Cell states and time are related, as earlier cell states transition into new cell states (such as during development). Bottom: Discrete cell states exhibit cell type–specific dynamics (such as circadian dynamics in mature cells). B: Simplified architecture of employed neural network. Count data is compressed and deconvolved into time–dependent and time–independent components. C: Top: Generation of synthetic per–cell time series by sampling from the time–independent latent space and then modifying the time–dependent component to project cells forward and backward in time. Bottom: Batch effect correction and imputation by constraining the second derivative of generated time series.

## 2 Results

To address the aforementioned challenges, we developed SNOW (SiNgle cell flOW map), an unsupervised probabilistic approach for the annotation, normalization, and analysis of single cell time series data. Briefly, SNOW implements a variational autoencoder [VAE] framework to simultaneously decompose the gene expression profile of each cell into time–dependent and time–independent components in a low dimensional latent space (Figure 1B). This design reflects the fact that while the gene expression of a mature cell (e.g., a neuron) may change in time (e.g., in response to a perturbation or circadian gene regulation), the cell type remains the same (i.e., a neuron remains a neuron). The time–independent component learned by the VAE thus encodes the identity of each cell, and is combined with a time–dependent factor to reconstruct the full gene expression profile. Importantly, the Universal Approximation Theorem [27, 28] implies that SNOW’s neural network architecture can articulate complex relationships between time and cellular identity contributing to gene expression without making any assumptions about their functional form, thereby directly addressing the situation depicted in Figure 1A where the dynamics depend on cell type.

As we will show, SNOW’s representation enables us to perform cell–type annotation using the time–independent latent space while also being able to study dynamics *without* requiring any cell type annotation. By changing the time coordinate in the latent space, the SNOW decoder can project individual cells forward and backward in time to generate cell–specific time–series that can be analyzed without the need to cluster or classify the cells (Figure 1C, top). By constraining the second derivative of the VAE output with respect to time, SNOW removes potential batch effects contaminating the time–series, enabling one to jointly perform batch integration and temporal analysis (which previously required two steps). We detail the SNOW method below and demonstrate how it can mitigate batch effects and capture biologically meaningful dynamics.

### 2.1 SNOW algorithm

We aim to achieve a number of things with SNOW. First, we wish to construct a time-independent characterization of the cell state to facilitate cell type annotation. This is achieved by minimizing the Wasserstein distance, a measure of distance between probability distributions, between the prior, *p*(*z*), and the latent distribution conditioned on sampling time, *q*(*z*|*t*). Second, we wish to map cells forward and backward in time such that the average of the model–generated gene expression time series across cells matches that of the population average (Figure 1C, top). To increase the smoothness of the interpolated trajectories, we incorporated in the loss function the second derivative of generated time series to penalize high curvature (see Methods for more detail). As a consequence of this second derivative loss, batch effects in the form of a sudden increase or decrease in expression can be detected and removed (Figure 1C, bottom). Third, we want to be able to correctly infer the sample collection time for an untimed sample, which is an active field of research in chronobiology [29–31]. To do this, we incorporated two additional terms in the loss function: one related to predicting the actual sampling time of each cell, and another related to predicting the sampling time of a cell after being mapped to another time by the model.

To achieve this, SNOW models the expression *x*_*gc*_ of gene *g* in cell *c* s a sample from a zero-inflated negative binomial (ZINB) distribution *P* (*x*_*gc*_|*l*_*c*_, *z*_*c*_, *t*) that depends on the observed library size of the cell (*l*_*c*_), the cell state (*z*_*c*_), and the time (*t*) of the observation. The cell state *z*_*c*_ is a low–dimensional vector computed by an encoder network that represents the *time–independent* biological variation contributing to *x*. To remove the effect of time in constructing the latent representation *z*_*c*_, we constrain the variational posterior conditioned on time *q*(*z*|*t*) to be close to the prior 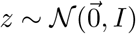.

The resulting time–independent latent space *z* of cell states has a number of appealing uses. It may, if desired, be used to conduct cell type annotation (Figure 1B). By changing *t* as an input to the decoder while holding the time–independent representation constant, we can generate a gene expression profile of a cell as it might appear at past or future times. In other words, we create an object similar to a flow map, in which the expected expression of the past/future state of any individual cell can be generated.

Technical details of SNOW’s probabilistic framework and loss function are given in the Methods section.

### 2.2 SNOW constructs biologically meaningful latent spaces

Time can have a profound impact on single cell data when it contributes to gene expression together with cell state. To see if SNOW can identify time–invariant structures presented in data, we constructed toy datasets that are composed of both a rhythmic component (genes that contain a cell type–specific phase) and a flat component (genes that contain a cell type–specific mean expression), as shown in Figure 2A. Details of the data may be found in the Methods. We generated 1000 cells each with an assigned cell-type label and sampling time. The cell type label (which defines the phase of the rhythmic component) and the sampling time of each cell jointly define the expression level of the rhythmic component. From our simple toy dataset, we observed that the incorporation of the rhythmic component resulted in the generation of tiny clusters on the UMAP [32] plot (Figure 2B), wherein cells segregated by both type and time. As expected, when only the flat component is used, the UMAP plot clustered according to the cell type of each cell (Figure 2B).

**Figure 2.**
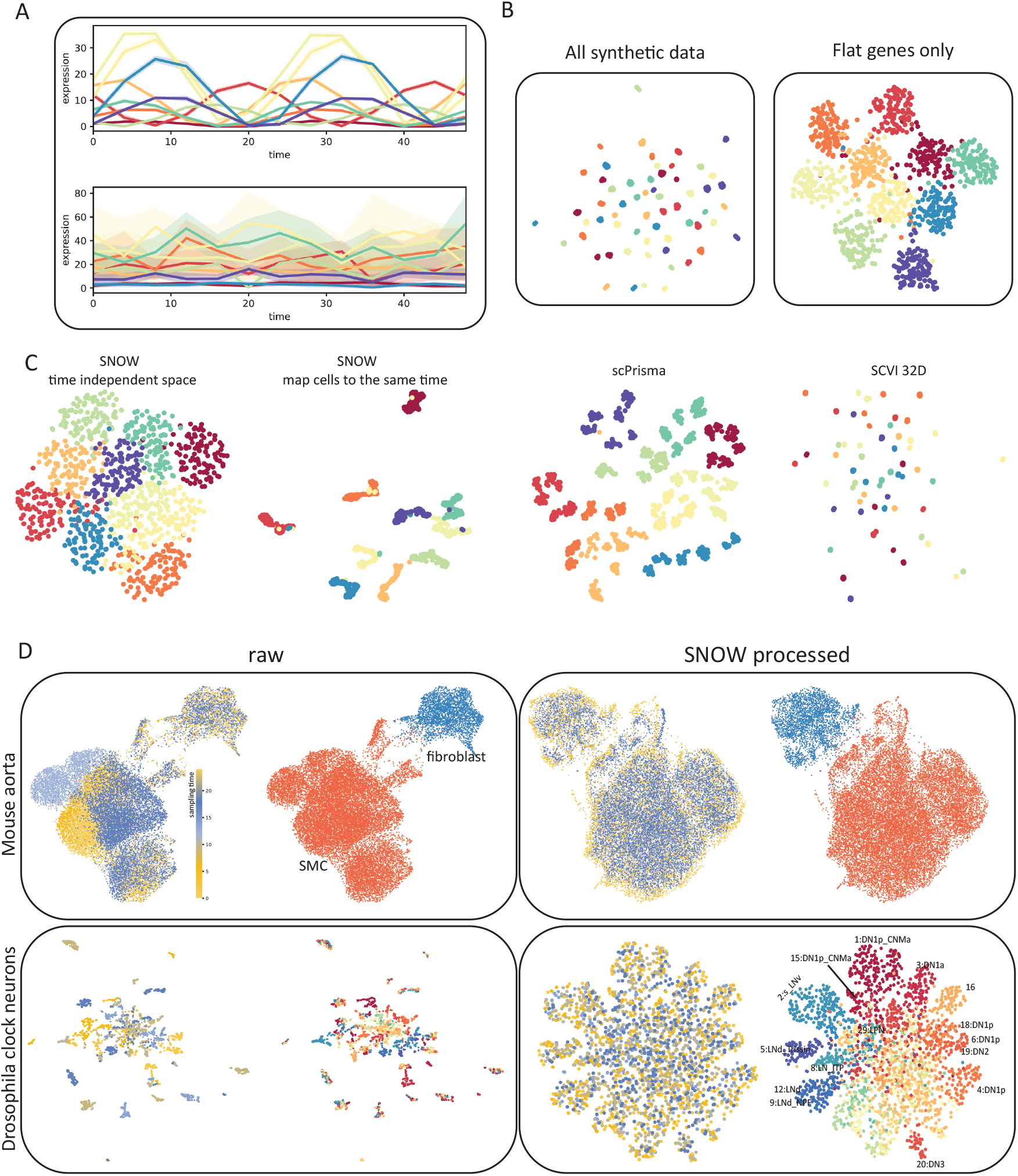
SNOW removes temporal effects while preserving biologically meaningful structure. A: Example toy data that consists of a rhythmic and a flat component, colors denote cell types. B: UMAP projection of the toy data using all genes and using only the flat component. C: Lowdimensional projections of the data generated by SNOW, scPrisma and scVI. Colors denote cell types D: Shown are UMAP plots of a mouse aorta dataset (top) and a drosophila neuron data dataset (bottom) using unprocessed (left) and SNOW-processed (right) data. Within each panel, the UMAP plots are colored according to annotated cell type (right image) and sampling time (left image). In the mouse aorta data without correction (top left), time separates the smooth muscle cell (SMC) cluster (orange) into subclusters. In the SNOW embedding of the same data (top right), the temporal effect has been removed and the SMCs and fibroblasts remain separated (top right). In the drosophila neuron dataset, UMAP shows clusters strongly dominated by time in the unprocessed data (bottom left), but by cell type in the processed data (bottom right). Cell type annotations by Ma et al. [10] are shown.

As described in the Methods, SNOW constructs a representation of the cell state that is independent of time and can be used for cell type annotation, either by using the latent space directly or by mapping cells to a common time. To test how SNOW performs, we applied it to the aforementioned toy dataset and compared it to scPrisma [33] and scVI [34]. As described in the methods, scPrisma [33] is used to decouple and filter the rhythmic component, while scVI [34] was used to treat the sampling time as batch labels and remove its effect. Of the three methods, SNOW is the only one that consistently captured the cell type information independently of the temporal variation (Figure 2C and Figure S2). Both scPrisma and scVI split the cell types into additional clusters based on time (Figure 2C). We further evaluated the performance of these three algorithms when the cell type dependence was only in the phase, only in the amplitude, or both (Figure S2A). We observed that scPrisma and scVI performed best when we have a cell type dependent amplitude but a cell type *independent* phase (Figure S2A). When the phase varies in a cell type specific manner, either alone or in conjunction with amplitude, scPrisma and scVI’s performance degraded. By contrast, SNOW had near–perfect performance in all cases (Figure S2A).

We also constructed a toy data that contained only one single cell type (Figure S2B), for which we expect a single cluster once the effect of time is removed. Even in this situation, we observed that neither scPrisma nor scVI could completely remove the temporal effect from the data, and cells continue to cluster by time in the UMAP space (Figure S2B). This observation suggests that the direct enforcement of time–independence imposed by SNOW can remove temporal effects that other methods cannot.

To illustrate the complexity introduced by time in real scRNA-seq datasets, we used UMAP to create lower dimensional embeddings of time–series scRNA-seq data collected from the drosophila clock neurons [10] and the mouse aorta [35], both with existing cell type annotations. We observed that the effect of time strongly drove clustering in the UMAP space (Figure 2D, left column). As illustrated in the top row of Figure 2D, while the UMAP space can separate the smooth muscle cells (SMCs) and fibroblasts, the SMC cluster contains subclusters, each corresponding to different sampling times. This effect is even stronger in the drosophila clock neurons, where the UMAP projection separates into small, disjoint clusters comprising cells sampled at particular points in time, often with mixed cell types (Figure 2D, second row).

We applied SNOW to both the drosophila and the mouse dataset and observed the successful removal of the temporal effect (Figure 2D, right column). Close examination of the SNOW latent space generated from the drosophila data (Figure 2D, bottom) reveals that we have retained variation attributable to cell type. Adding the original cell-type annotations to the UMAP plot of the SNOW–processed data (Figure 2D, bottom right), we find dorsal neurons (‘DN’s) located on the top and right side, and lateral neurons (‘LN’s) on the left (Figure 2D).

We next compared SNOW to scPrisma and scVI on the drosophila dataset. In contrast to the toy datasets, we observed that scPrisma performed worse than scVI in the real datasets (Figure S3B and C), where it removed information regarding cellular identity (Figure S3B). SNOW continued to outperform both. As a further example, we also applied SNOW to a time series dataset charting the regeneration of mouse lungs subjected to bleomycin-mediated injury [36] (Figure S5) and observed that cells significantly affected by bleomycin in the original gene expression space were embedded more closely to their untreated counterpart in the UMAP space generated from SNOW, revealing their common transcriptomic background (Figure S6). To then examine how bleomycin effects these cell types, one can simply combine the time dependent and time independent component to project the data back to the original gene expression space. In comparison, when we applied scVI to the same dataset, its latent space contained four clusters of cells (Figure S5B).

Finally, we tested whether trajectory inference methods such as Monocle [37–39] could be used to articulate the temporal variation. It should be noted that trajectory inference is a fundamentally different problem than that which we are trying to solve here. In trajectory inference, one assumes that the cells observed at a given time–point may be thought of as observations along a pseudotemporal trajectory, in which cells transition smoothly from one type to another. There is thus the assumption that transcriptomically similar cells are closer in time as the trajectory evolves from one cell to the next. In SNOW, we make the assumption that we are observing mature cells where the dynamics of gene expression in those cells depends both on time (which we observe directly by taking multiple samples over time) and cell type, without making the assumption that cells that are transcriptomically similar cells are closer in time.

Nevertheless, one might expect that a trajectory inference method would place cells of the same type from adjacent time–points closer to one another than cells of a different type, thereby articulating the trajectory of cells by organizing cells of the same type together and correctly ordering them in time. In the drosophila dataset, we would expect that these methods can first identify the fact that different cell types are traversing different trajectories, and for each cell type, identify the fact that cells are traveling along a circular trajectory. However, as illustrated in Figure S4A, time and cell type jointly affect the transcriptome. Without using algorithms like SNOW to disentangle this joint effect, the trajectory inferred by Monocle captured neither cell types nor the fact that cells are traversing a circular path. Using “batch corrected” data also does not solve the problem. In Figure S4B, we conducted trajectory inference with Monocle using its built–in method to conduct batch correction. While this space is now more consistent with cell types, the circular paths cells should take have been destroyed during batch correction, since batches and time are confounded. As a consequence, no substantial trajectories are inferred by Monocle.

### 2.3 SNOW maps cell forward and backward in time

SNOW generates a latent space that is independent of time and contains a decoder that reconstructs the transcriptome when the latent state and time are both supplied. In principle, then, it is possible to provide the latent representation of a samples cell with an *unsampled* time to generate an expression profile of that specific cell at another time–point. To test whether we can produce expression dynamics for each cell that is consistent with the population averages of its cell type, we generated *de novo* time series by concatenating the latent representation of a cell, *z*, with time, *t*. Because the concatenated *t* can differ from the sampling time of the cell, we refer to this as the “pseudo” sampling time. We generated time series using latent representations of the mouse aorta, which is sampled every six hours for one day, by using 100 equally spaced pseudo sampling times. One might then reasonably ask: if the generated data had in fact been observed data, would the encoder network have correctly identified the time that was used to generate the pseudo sample? By supplying the generated expression profile back to the encoder network, we observed that we are capable of re-inferring the pseudo sampling time of each cell accurately (Figure S7A), with a mean absolute error 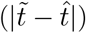 of 0.80 and 0.79 hours for the smooth muscle cells and the fibroblasts respectively (Figure S7B). Overlaying the mean absolute error on its UMAP projection identified no regions with particularly large errors (Figure S7C).

To further validate our approach, we averaged the generated time series for all cells from the same cell type and compared this population average to the experimental data (black lines in Figure 3). Using the well–characterized circadian genes as examples, we observed that while the generated expression time series for both the fibroblast and SMC show a considerable amount of diversity, the population average exhibits clear oscillatory dynamics and matches closely with empirical observation (Figure 3A, B). It is worth noting that no constraint was imposed during the training process to shape the generated population average. This observation suggests that the agreement between the observed and the generated dynamics is consequent of a successful deconstruction of the gene expression into time–dependent and time–independent components.

**Figure 3.**
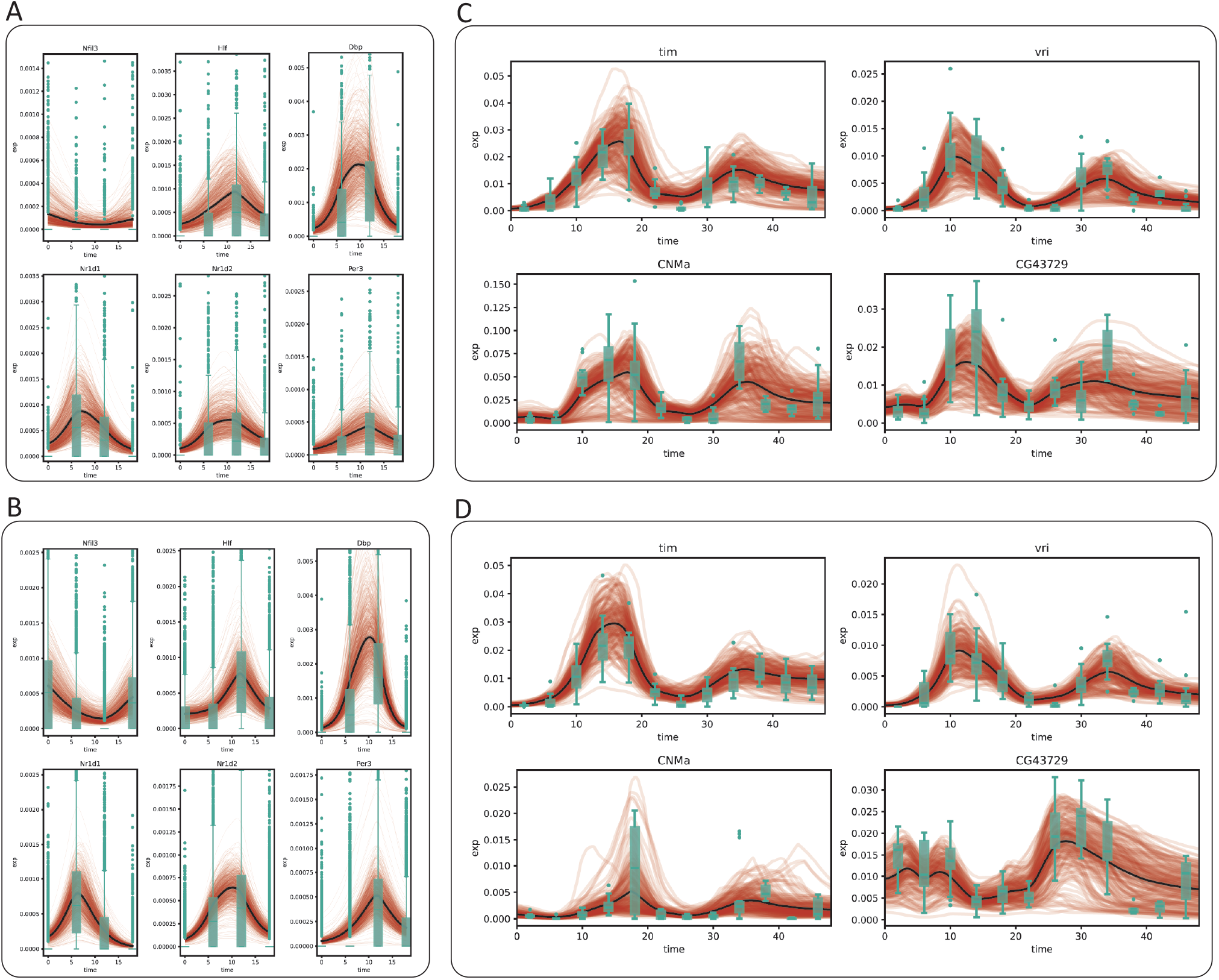
SNOW generates cell–level expression time series. Model generated cell-level expression time series (red), average of model generated time series (black) and observed gene expression (green boxplots) for the mouse aorta (A: mouse fibroblast), B: mouse SMC) and drosophila clock neuron (C: dorsal neurons, D: lateral neurons) datasets.

We next repeated this test on the clock neuron dataset, sampled every 4 hours for two days. As a sanity check, we tested whether SNOW’s encoder would recover the pseudo sampling time when the data generated by the decoder was fed back into it. We observed that our model remained competent at recovering the pseudo sampling times (Figure S8A), with an mean absolute error ranging from 1.5 hours to less than 3 hours. As before, we observed SNOW–generated oscillations in known circadian markers in concordance with experimental observation (Figure 3C, D). Despite the proximity of the 1:DN1p_CNMa cluster and the 2:s_LNv cluster in the UMAP space (Figure 2D, bottom left), we observed the mean expression level of the generated expression time series of *CNMa* to differ by ten fold, suggesting our usage of a fixed latent space standard deviation did not prevent the model from learning the dynamics particular to each cell type.

Interestingly, we found that the quality of the generated time series is tied to the size of the latent standard deviation (*σ*_*z*_). In the clock neuron dataset, we observed that small *σ*_*z*_ leads to damped oscillation in the long run (Figure S8B). However, this effect is not apparent in the mouse heart data (Figure S8B), potentially because of its larger sample size, simpler cell type composition, and fewer sampling times.

### 2.4 SNOW corrects batch effects

While batch effects can be difficult to identify and correct, the fact that samples are related in time provides a potential route of correction by prohibiting abrupt changes of expression, formally achieved by constraining the second derivative of the generated time series. To test whether SNOW can reduce the impact of batch effects in time–course data, we first identified genes that have been potentially affected. We consider a gene to be severely impacted by a batch effect if it is mostly detected only at a single time point. For those that are consistently detected across time points, we assume they are affected if their expression level at a particular time point is much higher than that of the rest (see Methods). With these two criteria, we identified 148 genes within the 1:DN1p_CNMa cluster from the clock neuron data and observed that 117 of them are considered to be features by Seurat [40]. As Seurat identifies features by looking for outliers on a mean–variance plot, it is expected, and alarming, that genes satisfying our criteria will be considered as features. By constructing time series using all sampled 1:DN1p_CNMa cells to span the entirety of the experiment, we observed that the generated signal is unaffected by the outlier samples (Figure 4A).

**Figure 4.**
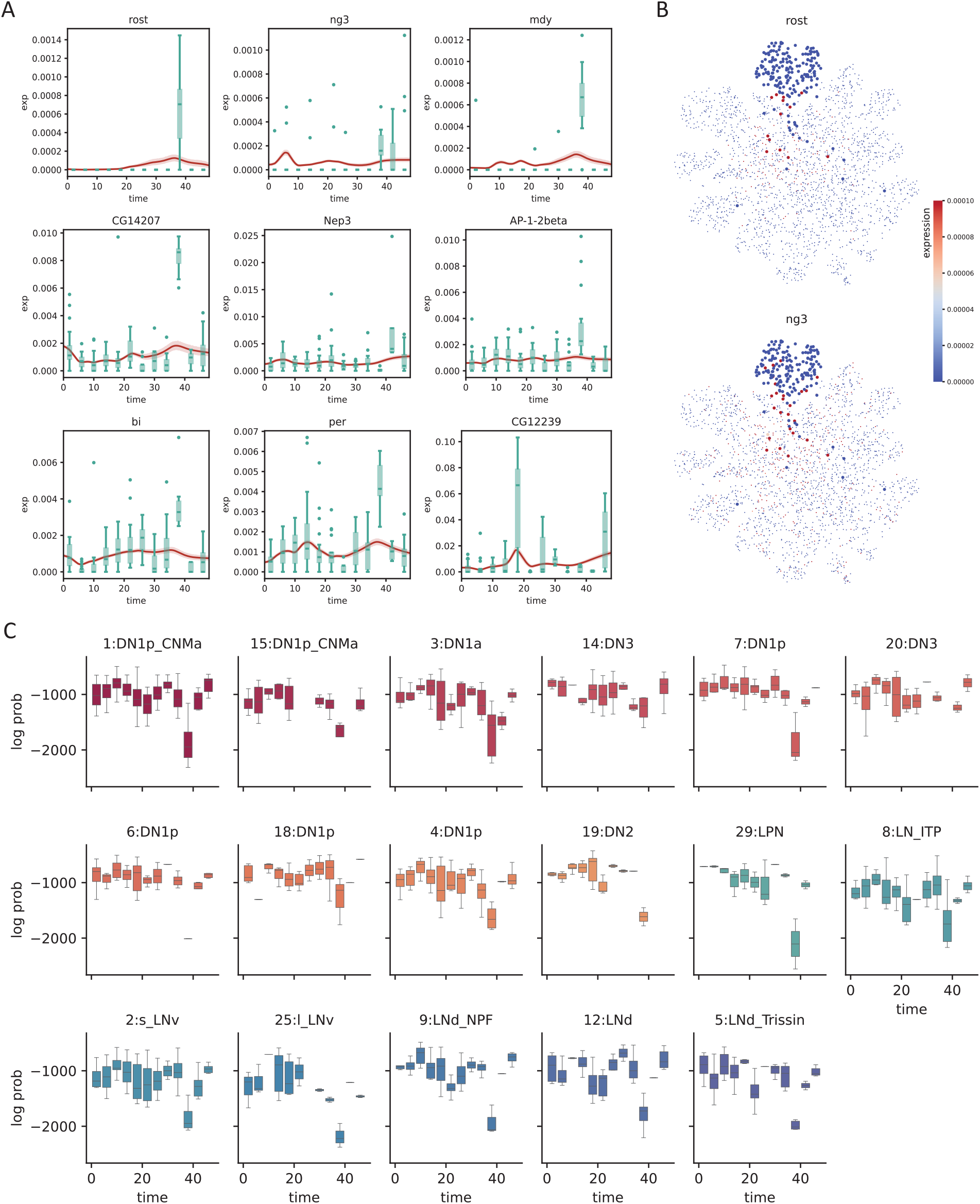
SNOW corrects potential batch effects. A: Examples showing SNOW–generated time series (average across cells shown as red lines, with shaded 95%CI) and the experimental observation (green boxplots). B: Gene expression of batch–affected genes overlayed on top of the UMAP projection of the clock neuron dataset. Cells belonging to the 1:DN1p_CNMa neuron cluster are plotted to be bigger. C: Box plots showing the log probability of observing each cell for the named clusters at each time–point.

Interestingly, we observed that a large proportion of the selected genes appear to be impacted by a batch effect at time ZT38. Direct visualization of the expression level of putative batch–affected genes on the UMAP space implies that these genes, which were not originally used as features, may contribute to the disagreement between the original cell type assignment and our latent space. For example, Figure 4B illustrates that cells annotated as 1:DN1p_CNMa neurons that had an elevated expression of batch–affected genes are located away from the main cluster. This suggests that what appears to be batch effect could be an artifact of cell type annotation. Since we can compute the likelihood of making an observation, if cells considered to be 1:DN1p_CNMa neurons at ZT38 were, in fact, of some other origin, cells collected at ZT38 would stand out from the rest of the time series, but the log–likelihood would not. To test this, we computed the log likelihoods of observing the experimental data and observed that gene-wise log likelihood also shows a sharp drop at the time when gene expression peaks (Figure S9), indicating that the observed expression level has a low probability of occurrence under our statistical model. Computing the log likelihood of observing the entire cell by summing up the probabilities of observing each gene, we noticed a drop at ZT38 for almost all cell types (Figure 4C, S9C), in agreement with our observation that a large fraction of the identified genes were impacted at ZT38. Additionally, this drop of log likelihood at ZT38 remained even when all cells were pooled together (Figure S9B), suggesting that the expression peaks we observed at ZT38 cannot solely be attributed to cell type assignment.

To summarize, we showed that SNOW can generate time series that are unaffected by outlier samples and that our underlying statistical framework is capable of detecting batch–affected genes.

### 2.5 SNOW allows unsupervised identification of circadian rhythms in gene expression

The discovery of tissue–specific circadian regulation [41] and advances in single cell technologies have led to studies that report cell-type specific circadian oscillation [10, 11]. While circadian time series conducted on the tissue level can be directly supplied to a number of readily available cycling detection algorithms [42, 43], single cell data requires some special considerations. First, proper cell type annotation requires the removal of all temporal effects. While this can be achieved via data integration, integrated data cannot be used for cycling detection, forcing users to conduct cell type annotation with integrated data but perform cycling detection with “raw” data. Moreover, one needs to choose whether to consider each cell as a replicate or to construct pseudobulk data for each time point. However, considering cells as replicates can be highly computationally inefficient, and it has been shown that constructing pseudobulk profiles can generate false positives, especially for genes with low expression [44].

With SNOW, we can generate a transcriptome-wide time series for each cell by projecting them forward and backward in time, thus enabling us to conduct cycling detection at the single cell level and detect fine expression differences between cell types (Figure S10). For each cell, we can obtain a *p*-value, phase estimate, and estimated amplitude for each gene via harmonic regression on its generated time-series. We note here that because these estimates are produced from generated data, care should be taken not to over–interpret the results; in particular, we recommend that the *p*-values from such an analysis be used *only* to rank/prioritize genes for follow up, rather than being considered in absolute terms. To test if the ordering of *p*-values obtained from the generated data reflect trends observed in the experimental data, we compared our results to the published list of per–cell–type cycling genes [10]. As demonstrated in Figure S11, genes that were reported to have rhythmic expression in multiple cell types had smaller average (across all cells) *p*-values and larger average (across all cells) amplitudes.

We then investigated the biological interpretation of the cell–level statistics. Unsurprisingly, known circadian genes *vri* and *tim* exhibited the strongest evidence of cycling (with the lowest– rankes average *p*-values across all cells). Overlaying harmonic regression *p*-values on the UMAP space showed that the core clock genes *vri, tim, Clk*, and *per* are highly cyclic in all cells (Figure 5A), as expected. We also overlayed the estimated oscillation amplitude and phase for each cell (Figure 5A). Interestingly, we observed high oscillation amplitudes of *vri* and *tim* in all labeled clusters. By contrast, cluster 16, an unnamed cluster, stood out for having a much lower amplitude despite its close proximity to the high–amplitude dorsal neurons on the UMAP space. Additionally, despite the fact that the phases of *vri, tim* and *Clk* were reported to be largely identical across cell types in the original (pseudobulk) analysis [10], we observed that SNOW is capable of discerning fine phase differences between clusters on a single cell level (Figure 5A).

**Figure 5.**
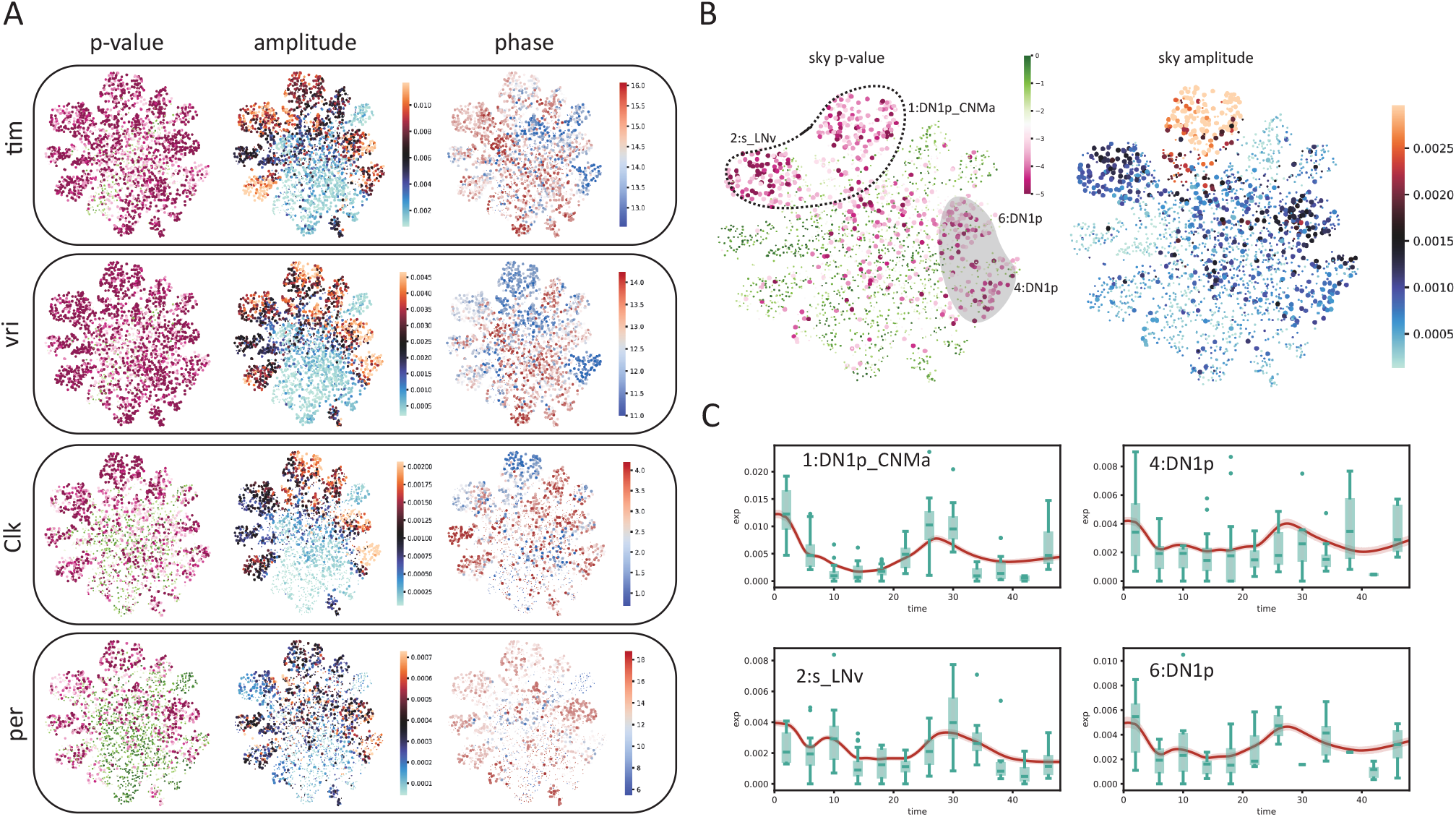
SNOW allows cycling detection at the single–cell level. A: Estimated *p*-values, amplitudes and phases (in hours) of known circadian genes (*tim, vri, Clk, per*) overlayed on the UMAP projection of the clock neuron dataset. Cells with *p* > 0.001 or amplitude smaller than 0.0001 were made small for better visualization. The *p*-value color scale is the same as panel B. B: Estimated *p*-values and amplitude of *sky*. The circled region indicates agreement between our analysis and that of Ma et al., and the shaded region indicates disagreements. C: SNOW–generated time series (average across cells shown as red lines with shaded 95%CI) and experimental observation (green boxes) of *sky* from cells within the circled and shaded region and panel B.

We also observed that there are cases where the harmonic regression *p*-values from flat genes are low, which leads to disagreement between our analysis and the published cycling genes. By looking at the estimated amplitudes, we found that these disagreements can be resolved by using amplitude criteria that exclude cells/clusters with low oscillation amplitudes (Figure S12). For example, *sky*, which was reported to be cycling in the 2:s_LNv and 1:DN1p_CNMa clusters, also appeared to be cycling in two other DN1p clusters (Figure 5B, left panel). While the estimated amplitudes of *sky* from the two DN1p clusters are smaller than that of the 1:DN1p_CNMa cluster, they are similar to that of 2:s_LNv neurons (Figure 5B, right panel). A closer look at the time series generated from the two DN1p groups revealed expression dynamics distinct from that of 1:DN1p_CNMa but similar to that of 2:s_LNv, suggesting *sky* may be cycling in a larger population of dorsal neurons than previously believed (Figure 5C). Another gene, *Ddc*, which was also reported to cycle in the 2:s_LNv neurons, showed high *p*-values and low amplitudes in our analysis (Figure S13A). Comparing SNOW–generated time series to the experimental observations (Figure S13B) suggests that this may have been a false positive in the original analysis. On the other hand, we observed that two dorsal neuron groups (7:DN1p, 20:DN3) in which *Ddc* was not reported to be cycling originally showed low *p*-values and high amplitudes in the SNOW generated data (Figure S13B), possibly suggesting a false negative (Figure S13B).

In summary, we showed that SNOW may be used to help the identification of rhythmic genes by first generating time series for each cell, and then conducting cycling detection on a single cell level. By doing this, cycling detection analysis does not depend on the accuracy of cell type annotation. This suggests that it can be used in combination with traditional analyses that first assign cell types prior to pseudobulking for cycling detection. For example, it can increase the confidence in the identification of cycling genes by confirming that they are rhythmic in the majority of individual cells; detect potential false negatives in the pseudobulk analysis (especially for rare cell types that may not be sampled at all time points); and avoid false-positives by removing potential batch effects. It can also potentially identify subsets of cells of a single type (or a single cluster) that are differentially cycling, an effect that may be missed in analysis where cells of the same type are treated as replicates or pseudobulked for cycling detection.

## 3 Discussion

We presented SNOW (SiNgle cell flOW map), a deep learning framework for the annotation, normalization, and simulation of single–cell time series scRNA-seq data. SNOW is designed for cases in which cell type does not change with time (i.e., no differentiation) but gene expression changes in a cell-type dependent way. Circadian transcriptional regulation is one example of this: it is estimated that nearly half of the genome is under circadian transcriptional control, with the identity of the cycling genes being specific to the tissue/cell type [41]. Over the course of the day, the cells change their gene expression, but they do not change their identity (a neuron remains a neuron, a fibroblast remains a fibroblast, etc.). Likewise, mature cells may exhibit time–dependent transcriptional changes in response to an external perturbation [22, 45].

SNOW uses a variational autoencoder to model gene expression as a zero-inflated negative distribution [34, 46] that depends on time-dependent and time-independent components. The time– independent latent coordinates can be used directly for cell type annotation, and the time–dependent component can be used to generate artificial time series for individual cells (in a manner similar to a flow map, but without the need or computationally expensive numerical integration). We demonstrated the utility of SNOW by applying it to multiple single–cell datasets with different cell numbers, sampling frequencies, and sequencing depths.

SNOW has a number of advantages. First, most methods for analyzing single cell time series data focus on developmental processes, in which one may assume that the temporal dynamics reflect changes in cell type. These methods largely rely on finding an optimal transport map between cells sampled at distinct time points [15, 17, 19, 20]. By contrast, SNOW is designed to elucidate gene expression dynamics in mature cells where expression changes with time in a cell type–dependent manner (Figure 1A). In such cases, it is desirable to deconvolve the contributions of cell–type and time. Previous methods have approached this problem with a two–step procedure in which temporal information is first removed for cell type identification and then reintroduced for temporal analysis [10, 22]. By contrast, SNOW addresses this issue by directly learning a latent representation that simultaneously articulates both the time–dependent and time–independent contributions to gene expression, enabling temporal analysis without the need to classify cells.

Second, we demonstrate that SNOW can be used to identify and eliminate batch effects. By modeling count data with a zero inflated negative binomial distribution, we were able to identify samples that are likely to be batch-affected by using the estimated probability of observing their gene expression profiles. Timepoints with particularly low probability across all cells, such as those shown in Figure 4C, are likely to be batch–affected outliers in the time–series and merit further investigation. In the case where a technical batch effect is suspected, corrected values can be estimated using SNOW’s decoder to generating gene expression observations for each cell at the affected timepoint.

Third, SNOW is capable of generating time–series data for individual cells, providing a novel avenue for single–cell time–series analysis. This can be used in a number of interesting ways. For example, previous work has shown that even seemingly simple tasks such as cycling detection (identifying genes with circadian oscillation) is non-trivial in single–cell data due to the fact that cells are destroyed at each timestep, requiring cells to be clustered and identified to make inferences across time [44]. Here we demonstrated that SNOW could be used to generate *per–cell* timeseries, mimicking what might happen if the cells could be observed continuously. Cycling detection (or other temporal analyses) can then be performed on an individual cell basis, generating inferences about which genes are cycling in which cells, and with what phases and amplitudes. These inferences may then be probed in followup experiments (e.g., using live cell imaging).

It should be noted, of course, that those “per-cell” inferences are only as good as the generated data, and thus should be treated of as “hypothesis generating” (and, in particular, any resulting *p*-values should only be used to prioritize genes based on their rank, rather than reported in absolute terms). The fact that the generated data recapitulates the observed expression data (as illustrated in Figures 3 and 5) suggests that inferences made from the generated data are not unreasonable. Why, then, might someone want to perform the analysis using simulated data rather than (or in addition to) real observations?

Consider the analysis protocol for the observed data, where the cells observed at each time– point are distinct cells (indeed, the number of cells may even change from one time–point to the next). In order to link observations at one timepoint to those in the next, cells are clustered or classified into types, and the behavior of the clusters over time is analyzed. Typically, cells within a cluster are averaged to yield a “pseudobulk” expression profile for each cell type at each timepoint (or, alternatively, treated as replicate observations of a particular cell–type), and the downstream analysis is performed at the cluster/cell-type level. The success of this analysis not only depends on the correctness of the clustering/classification, it also makes the assumption that cells of the same type have similar dynamics, i.e., that there is no heterogeneity within a particular cell type — an assumption that may not be correct.

By contrast, using the SNOW-generated per-cell data does not require the identification of cell types or any clustering. This avoids propagating an errors in cell type identification. It also means that we are able to detect heterogeneous dynamics within cell–types, rather than making the implicit assumption that all cells of a given type will behave similarly. This approach can be paired with traditional analyses to increase the confidence in detected cycling genes and potentially identify false positive/negative cyclers that can arise in the traditional two–stage analysis [44]. This capability can be useful in non-circadian contexts as well, such as when one wishes to identify genes that follow a specific pattern during differentiation or drug perturbation.

The ability of SNOW to generate cell–level trajectories also enables the generation of novel hypothesis that would not be possible otherwise. For instance, one could perform analysis shown in Figure 5 and then bring cell identity back into it to ask: do all cells of a particular type have the same dynamics (phase, amplitude), or is there heterogeneity for certain cell types? Other analyses are also possible. For example, SNOW allows one to identify genes that may mediate the amplitude of circadian oscillation by looking for genes with an expression level that correlates with oscillation amplitude of the core clock genes in each cell. Additionally, one could also imagine supplying SNOW-generated time series data to network reconstruction algorithms to further dissect the details of gene-gene interaction.

Several existing methods bear some similarities to SNOW, with important differences. scVI [34], DCA [46], and many other methods [47–50] are all built on variational autoencoders [18], which are typically trained via the optimization of the evidence lower bound (ELBO). However, as the ELBO only constrains the latent space via the KL divergence (Eq 6), it may generate correlated latent dimensions and fail to enforce the assumption that the prior distribution has identity covariance, enlarging the difference between ELBO and the actual log likelihood, log(*p*(*x*)). In this situation, one would fail to generate “realistic” virtual gene expression profiles by passing samples drawn from the prior distribution through the decoder network. While having an “irregular” latent space that fails to match the prior distribution may not impact the performance of the model in other tasks such as clustering and identifying cell types, enforcing independence between the latent dimensions is known to improve model interpretability [51, 52]. To address this, scNODE [17] and scVIS [47] introduced a scaling factor added to the KL divergence term (similar to *β*-VAEs [51]) to enforce a stronger constraint on the latent space, thereby encouraging a more efficient representation of the data. More recently, various methods have been proposed [52–55] to directly enforce independence between latent dimensions via minimizing *D*_*KL*_ (*q*(*z*)|| Π_*d*_ *q*(*z*_*d*_)). In SNOW, we enforced independence between latent dimensions and alignment with respect to the prior distribution simultaneously by minimizing the sliced Wasserstein distance.

As mentioned in previous sections, we developed SNOW to solve the following problem: in the situation where both cell state and time affects gene expression, removing temporal effects to facilitate cell type annotation also removes biologically meaningful gene expression dynamics. This problem is related to what MrVI [48] attempts to solve by constructing a sample-unaware representation (*u*) and a sample-aware representation (*z*), where *u* is used to conduct cell type annotation and *z* is used to model how sample related covariates (such as a batch or a time–point) affect gene expression. In some sense, SNOW and MrVI are designed to solve a common problem, except that SNOW specializes in continuous covariates (time) and MrVI in discrete covariates. Our explicit enforcement of statistical independence between the latent space and time, which is absent in both MrVI and scVI, naturally defines cell state as a time–invariant quantity. By supplying the decoder with time and the time–independent representation of cell type, SNOW can generate data “sampled” from intermediate time points, which cannot happen if time is simply treated as batch label, as it is in MrVI. SNOW also has the additional benefit of enforcing smoothness by constraining the second derivative with respect to time, which is not possible if time is treated as a categorical variable.

SNOW has a few limitations. Most significantly, we assume that the distribution of cell types that does not change with time, which allows us to construct the time–independent latent space by minimizing the sliced Wasserstein distance across all timepoints. In developmental systems, where the distribution changes as new cell types emerge or vanish, this assumption may be violated. We also acknowledge that SNOW does not consider the effect of growth, which is the focus of a number of existing approaches [15, 20, 26]. It has been shown that taking growth into consideration can improve RNA velocity estimation by removing false cell state transitions [26]. However, in the problem we seek to address, we reason that growth is less relevant because we are not examining dynamics generated by cell state transitions such as division. Rather, the dynamics that we study are intrinsic to each cell (such as the self-driven circadian oscillation of the drosophila clock neurons subtypes).

We expect our work to be of interest to those studying dynamic processes in complex tissues. To our knowledge, SNOW is the first method that provides simultaneous batch correction and time series analysis, which is typically done in two separate steps (as in [22]). This allows time– series analyses to be performed without clustering or classifying the cells. Our decomposition of gene expression allowed us to decipher datasets that seem to make little sense at a first glance (Figure 2D). Additional features can be easily added into our method to handle more complex datasets, and approaches employed in our work, such as data integration or the enforcement of statistical independence, can also be extracted and adopted for other analyses. In the future, we envision that our approach can be used to extract the core qualities of cells sampled from more complex conditions, such as those jointly influenced by age, gender, and even the disease stage of the donor. In these cases, identifying such core qualities may help us better predict drug response, patient outcomes and even help the design of improved dosing strategy.

## 4 Conclusion

We presented SNOW, a novel computational approach to characterize single cell time series data through the decomposition of gene expression into a time–dependent and a time–independent space. Our approach allows the unprecedented opportunity to conduct time series analysis on the level of single cells through the generation of cell–level data. SNOW can be used to deconvolve the effects of time and cell-type for better cell type identification; to detect and mitigate batch effects; to conduct temporal analysis without the need for cell clustering or classification; and to generate per–cell time series data that can lead to testable hypotheses. Used in conjuction with existing methods, SNOW can provide novel insights and serve as the basis for future methods development.

## 5 Methods

### 5.1 SNOW’s probabilistic framework

SNOW models the count matrix **X** ∈ ℝ^*C*×*G*^ with a zero-inflated, negative binomial (ZINB) distribution [34, 46], where *C* and *G* are the number of cells and genes in the sample, respectively. Without zero-inflation, a given entry within **X**, *X*_*cg*_, is modeled as:

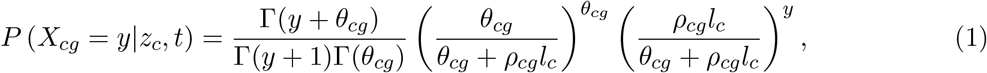

where Γ(*·*) is the standard gamma function, *z*_*c*_ the (time-independent) encoded state of **X**_*c*_ sampled at *t, θ*_*cg*_ the gene– and cell–specific inverse dispersion, *l*_*c*_ the library size of cell *c*, and *ρ*_*cg*_ the count fraction of gene *g* in cell *c* such that ∑_*i*_ *ρ*_*ci*_ = 1. *θ* and *ρ* are optimized using neural networks *f*_*θ*_ and *f*_*ρ*_ respectively.

Zero-inflation is added with the following form:

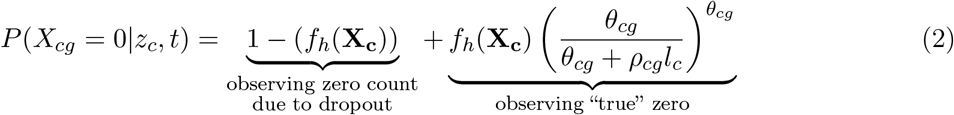

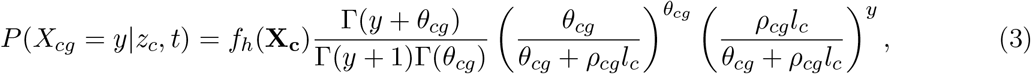

where *f*_*h*_(*·*) is parameterized with a neural network. Since elements of **X**_*c*_ ∈ ℝ^*G*^ are conditionally independent of each other given *z* and *t* (*∀i* ≠ *j, P* (*X*_*ci*_|*z, t, X*_*cj*_) = *P* (*X*_*ci*_|*z, t*)), we can compute the probability of observing the count profile of a particular cell as:

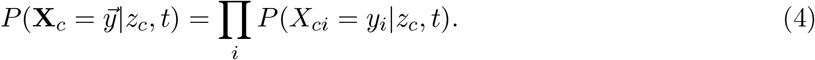

Or equivalently:

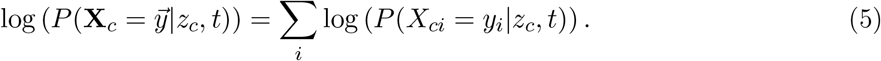

Our framework allows the generation of “virtual” cells by assuming a Gaussian prior, a commonly used prior for building variational auto-encoders, as follows:

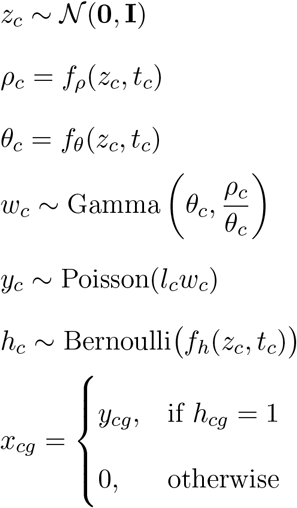

where *z*_*c*_ is the time-independent latent representation of a cell; *ρ*_*c*_ ∈ [0, 1] is the normalized expression profile (or count fraction) enforced by using a softmax activation function in the last layer of *f*_*ρ*_; *x*_*c*_ ∈ℕ^*G*^ is the count profile of the virtual cell; and *l*_*c*_ is the observed the library size. The Gamma-Poisson process generates *y*_*c*_ ∈ℕ^*G*^ following a negative binomial distribution with mean *ρ*_*c*_*l*_*c*_, while *h*_*c*_ is a binary vector that represents dropouts. *f*_*ρ*_, *f*_*θ*_, and *f*_*h*_ are neural networks that map the latent space and time back to the full gene space, 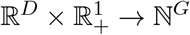.

### 5.2 SNOW loss function

A number of methods have used variational autoencoders (VAEs) [18] to model count data from single-cell RNA seq [17, 34, 47]. All have used loss functions reminiscent of the evidence lower bound (ELBO), which constrains the shape of the latent space *q*(*z*) indirectly via the KL-divergence term:

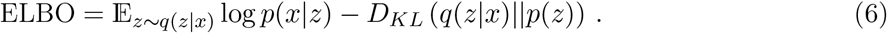

(See derivation of ELBO in Supplement.) In the above expression, *p*(*z*), the prior distribution of the representations *z*, has been chosen for convenience to be 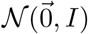 and *q*(*z*) is the variational posterior distribution of *z* constructed by the encoder network. The KL-divergence term provides the model some level of robustness, as it essentially requires points near *z* in the latent space to be decoded into similar objects. However, as the dimensionality of the data grows, the log-likelihood term of the ELBO will dominate over the regularizing KL-divergence term. While this is unaccounted for in scVI [34], both scNODE [17] and scVIS [47] incorporate scaling factors to maintain the strength of the regularization of the latent space. By definition, maximizing ELBO leads to the maximization of the marginal log likelihood (log *p*(*x*)),

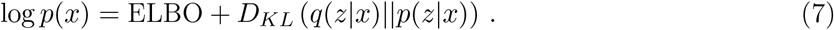

When *D*_*KL*_ (*q*(*z*|*x*)||*p*(*z*|*x*)) = 0, or equivalently *q*(*z*|*x*) = *p*(*z*|*x*), the ELBO will be equal to the marginal log likelihood of *x* and *p*(*z*) = ∫ *q*(*z*|*x*)*p*(*x*)*dx* = *q*(*z*). However, when the ELBO is not tight, its optimization can lead to an enlargement of the approximation error, *D*_*KL*_(*p*(*z*|*x*)||(*q*(*z*|*x*)). To account for this, SNOW regularizes the latent space directly by minimizing the distance between the latent distribution (*q*(*z*)) and the prior (*p*(*z*)) as measured by the Wasserstein distance. Briefly, in addition to the log likelihood term, the SNOW loss function begins with two main regularization terms, the former of which regularizes the latent space and the latter of which enables predictions of the sampling time:

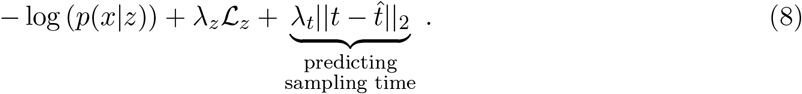

In the above expression, ℒ_*z*_ regularizes the latent space *q*(*z*) and enforces time–independence via:

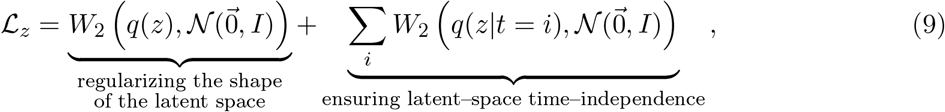

where *W*_2_(*q, p*) denotes the Wasserstein-2 distance between distributions *p* and *q*. This regularization enables the generation of a “virtual” cell when *z* is sampled from 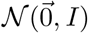. To ensure our model can generate proper “synthetic” cells sampled from different time points, we enforced two things. First, the time–independent components of the “synthetic” cells should follow the same distribution as that of the real cells (a Gaussian distribution). Second, the sampling time of the “synthetic” cells should remain predictable. To achieve this, we therefore impose:

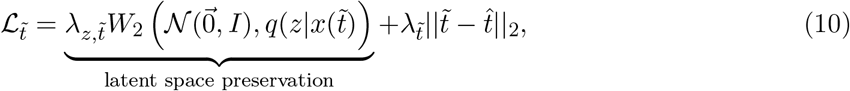

where 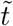 is the sampling time of the “synthetic” cells. And finally, we constrain the second derivative of the generated time series to enforce smoothness:

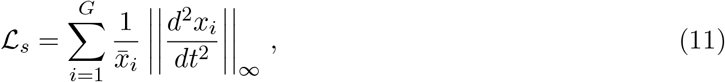

where *G* is the number of genes and 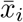 is the average of *x*_*i*_ over all generated time points. In practice, we find that computing ℒ_*s*_ for a randomly selected gene, *r*, in each training loop to be computationally cheaper and sufficient to generate smooth time series, giving the final form of our loss function:

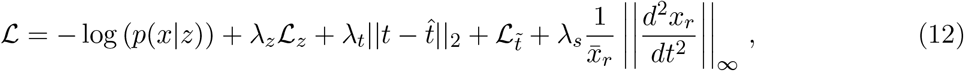

which preserves the latent space distribution, its time independence, and ensures the smoothness of the generated time series.

In practice, we simplify the calculation by replacing the Wasserstein-2 distance *W*_2_ with a more computationally tractable form, the sliced Wasserstein distance (*Ŵ* _2_) [56], defined as:

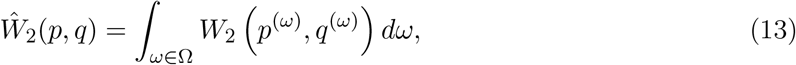

where the distributions *p*^(*ω*)^ and *q*^(*ω*)^ can be generated by first sampling from *p* and *q* directly before projecting them in a random direction, *ω*, sampled uniformly from the unit sphere 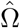. Given a set of data points 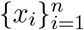 with an unknown underlying distribution *q*(*x*), the sliced-Wasserstein distance with respect to a known distribution, such as the standard normal, can be easily computed as:

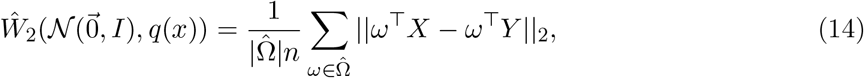

where 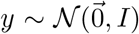 and we assume the columns of *X* and *Y* are sorted such that elements of both *ω*^*⊤*^*X* and *ω*^*⊤*^*Y* are arranged in ascending/descending order.

### 5.3 Neural network optimization

By default, SNOW uses a 3–layer encoder neural network with 256 fully connected neurons per layer and ReLU activation to project count data onto a 32 dimensional latent space (*z*_*c*_). Subsequently, *z*_*c*_ and *t* were used as input to individual neural networks (*f*_*ρ*_, *f*_*θ*_ and *f*_*h*_) with the same structure as the encoder network to generate the count fraction, inverse dispersion and dropout probability. To ensure that *f*_*h*_ generates probabilities, its last layer is activated by a sigmoid function so that its output ranges from 0 to 1. We further clamped the dropout probability between 0.01 and 0.99 to prevent the appearance of log(0). As mentioned above, the last layer of *f*_*ρ*_ is activated by a softmax function to enforce the sum of its output. During each training loop, we focus only on a randomly selected small subset of the data, by default 300. Everything within the loss function is computed from information contained within this subset of 300 cells, which enables our method to be applied to larger datasets in a memory–efficient manner.

In all test cases, the optimization of the model parameters was done with the ADAM [57] optimizer as implemented by pytorch [58] with a learning rate of 0.0005, *β*_1_ = 0.8, *β*_2_ = 0.9, and a weight decay of 0.0001. No scheduler was used to change the learning rate during the training process.

### 5.4 Construction of toy datasets

To test SNOW with data where the dynamics are known, we constructed a “toy” dataset containing two types of genes: “flat” genes and “rhythmic” genes. Each cell is randomly assigned a sampling time, *t*, from [0, 4, 8, 12, 16, 20, 24, 28, 32, 36, 40, 44, 48]. We simulated the expression of 1000 cells. In Figure 2, we simulated 120 rhythmic genes and 30 flat genes. In Figure S2, we simulated 50 rhythmic genes and 100 flat genes. For the *i*^th^ flat gene in cell type *j*, its expression level at time *t*,*x*_*i*_(*t*), is defined as:

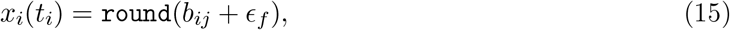

where *ϵ* is added noise, *b*_*ij*_ is the basal expression of the flat gene *i* in cell type *j* and `round`(*·*) maps its input to the nearest integer. For the *i*^th^ rhythmic gene in cell type *j*, its expression level at time *t* is defined as:

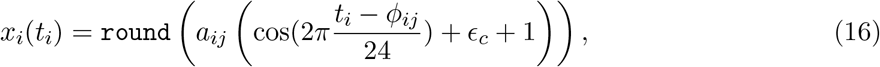

where *a*_*ij*_, *ϕ*_*ij*_ is the amplitude and phase of gene *i* in cell type *j*, and the +1 term lifts the cosine wave to have a zero minimum. Amplitudes *a*_*ij*_ and baseline expressions *b*_*ij*_ were chosen uniformly at random on [0, 20], and phases *ϕ*_*ij*_ were chosen uniformly at random on [0, 2*π*). Noise levels for the flat and cycling genes were drawn on *ϵ*_*f*_ ∼𝒩 (0, 2) and *ϵ*_*c*_ ∼𝒩 (0, 0.1), respectively. In Figure 2, *a*_*ij*_ is only gene dependent thus it can be reduced to *a*_*i*_. In Figure S2A, we considered the situation when only *a*, only *ϕ*, or both *a* and *ϕ* are cell type dependent. Negative values are thresholded to 0.

### 5.5 Application datasets

We applied SNOW to three sc-RNAseq datasets, including a circadian drosophila clock neuron dataset, a circadian mouse aorta dataset, and a lung regeneration dataset.

#### The circadian drosophila clock neuron dataset

The drosophila clock neuron dataset [10] (mean UMI/cell = 20060) was collected from *Drosophila* clock neurons every four hours with two replicates (12 time points in total) under both light-dark (LD) and dark-dark (DD) cycles. We focused our analysis on cells subject to the LD cycle, which contains 2325 cells. Count data was downloaded from the Gene Expression Omnibus under the accession code GSE157504 and the relevant metadata from https://github.com/rosbashlab/scRNA_seq_clock_neurons. Data integration was conducted using the IntegrateData function from Seurat [23] with ndim = 1:50, and k.weight=100. The resulting counts were used as input to the model.

#### The circadian mouse aorta dataset

The mouse aorta dataset [35] (mean UMI/cell = 14181) was collected every 6 hours (4 time points in total) under LD conditions, with a total of 21998 cells. H5ad files of the smooth muscle cells (SMC) and fibroblasts were downloaded from https://www.dropbox.com/sh/tl0ty163vyg265i/AAApt14eybExMMPK7VVDmfvga. Raw counts were used as input to the model.

#### The lung regeneration dataset

The lung regeneration dataset [36] (mean UMI/cell = 1585) was collected every day for two weeks (day 1 through day 14), and on day 21, 28, 36 and 54. We used AT2 cells, cilliated cells and club cells because they are activated after bleomycin treatment, resulting in a total of 24383 cells. Gene expression data were downloaded from https://www.ncbi.nlm.nih.gov/geo/query/acc.cgi?acc=GSE141259 along with the associated metadata. Raw counts were used as input to the model.

### 5.6 Application of compared algorithms

**scVI [34]** scVI is implemented following https://docs.scvi-tools.org/en/stable/tutorials/notebooks/scrna/harmonization.html. Raw count data is used as input to scVI, and the sampling time of each cell is used as the batch key. We chose a ZINB to model gene expression and set network hidden unit to 256 and the dimension of the latent space to be 32 to match that of SNOW.

**scPrisma [33]** scPrisma is implemented following https://github.com/nitzanlab/scPrisma/blob/master/tutorials/cpu/tutorial_prior_knowledge_linear_and_cyclic_cpu.ipynb using the `filtering_cyclic_torch` function. The resulting filtering matrix **F**, which has the same dimension as the data matrix (**X**), is used to filter the data matrix by element-wise multiplication.

**Monocle [37–39]** Monocle3 is implemented following https://cole-trapnell-lab.github.io/monocle3/docs/trajectories/ using default parameters.

### 5.7 Criteria for identifying batch effects

To identify genes potentially affected by batch effects, we looked for two types of patterns: spurious expression and spurious detection. We consider a gene to have spurious expression if its maximum normalized expression at one time point is five times higher than its average over all time points; and we consider a gene to have spurious detection if its maximum capture rate (number of cells that contain a said gene over the total number of cells collected at this time point) at one time point is five times greater than its mean capture rate over all time points. Here, we used the empirical *ρ* as normalized expression. To exclude genes that are almost never detected, we only used those with an average normalized expression (across all time points) greater than 0.00001 and an average capture rate (across all time points) over 5%. In the clock neuron dataset, this analysis identifies 124 genes with unusual capture rates and 24 genes with unusual expression, with zero intersection.

### 5.8 Detecting circadian behavior on a single cell level

To conduct cycling detection for each individual cell, we generated pseudo samples for each cell by concatenating the time independent representation of the cell state, *z*, with time, *t*. Using the SNOW decoder, we generated time series comprising 24 time points spanning 48 hours. We then conducted harmonic regression on these time series, resulting in a *p* value, a phase estimate, and an amplitude estimate for each gene from each cell.

## 6 Acknowledgements

This work was supported by NSF grant DMS-1764421, Simons Foundation grant 597491, and NIH grant R01AG068579.

## 7 Code and Data Availability

Code for our analysis is available on bitbucket (https://bitbucket.org/biocomplexity/snow/src/main/).

