## supplementary figures for "Variational inference of single cell time series"

### 8 Supplemental material

#### 8.1 Deriving the variational lower bound

The count matrix $\mathbf{X}$ follows a complicated distribution $p(x)$, which is difficult to sample from. We aim to sample from a simpler distribution $p(z)$, found by maximizing $p(x)$ or the evidence lower bound (ELBO):

$$\begin{aligned}
\log p(x) &= \log \int_z p(x,z)dz \\
&= \log \int_z p(x,z)\frac{q(z|x)}{q(z|x)}dz \\
&= \log \mathbb{E}_{z\sim q(z|x)}\left[\frac{p(x,z)}{q(z|x)}\right] && \text{(A-1)} \\
&\geq \underbrace{\mathbb{E}_{z\sim q(z|x)}\left[\log\frac{p(x,z)}{q(z|x)}\right]}_{\text{evidence lower bound}}, && \text{(A-2)}
\end{aligned}$$

where the inequality is a consequence of Jensen's inequality. In practice, the variational posterior $q(z|x)$, is chosen such that it is easy to sample from. To quantify how it differs from the true posterior $p(z|x)$, one may compute the Kullback-Leibler divergence:

$$\begin{aligned}
D_{KL}\big(q(z|x)||p(z|x)\big) &= -\int_z q(z|x)\log\frac{p(z|x)}{q(z|x)}dz \\
&= -\int_z q(z|x)\log\frac{p(z,x)}{q(z|x)p(x)}dz \\
&= -\text{ELBO} + \log p(x). && \text{(A-3)}
\end{aligned}$$

Rearranging, we obtain:

$$\log p(x) = \text{ELBO} + D_{KL}\big(q(z|x)||p(z|x)\big). \qquad \text{(A-4)}$$

Additionally, the ELBO can be further expanded:

$$\begin{aligned}\text{ELBO} &= \int_z q(z|x)\log\frac{p(x,z)}{q(z|x)}dz \\ &= \int_z q(z|x)\log\frac{p(x|z)p(z)}{q(z|x)}dz \\ &= \int_z q(z|x)\log p(x|z)dz + \int_z q(z|x)\log\frac{p(z)}{q(z|x)}dz \\ &= \mathbb{E}_{z\sim q(z|x)}\big(\log p(x|z)\big) - D_{KL}\big(q(z|x)||p(z)\big). \end{aligned} \tag{A-5}$$

While KL divergence will be difficult to compute in most cases, closed form exists if $q(z|x) \sim \mathcal{N}(\mu_1, \Sigma_1)$ and $p(z) \sim \mathcal{N}(\mu_2, \Sigma_2)$:

$$D_{KL}\big(q(z|x)||p(z)\big) = \frac{1}{2}\left(\log\frac{|\Sigma_2|}{|\Sigma_1|} - n + \text{Tr}(\Sigma_2^{-1}\Sigma_1) + (\mu_2-\mu_1)^\top\Sigma_2^{-1}(\mu_2-\mu_1)\right) \tag{A-6}$$

By assuming $\mu_2 = \vec{0}$ and $\Sigma_2 = I$, the above expression can be further simplified to:

$$\begin{aligned} D_{KL}\big(q(z|x)||p(z)\big) &= \frac{1}{2}\left(\log\frac{|I|}{|\Sigma_1|} - n + \text{Tr}(I^{-1}\Sigma_1) + (\vec{0}-\mu_1)^\top I^{-1}(\vec{0}-\mu_1)\right) \\ &= \frac{1}{2}\left(-\log|\Sigma_1| - n + \text{Tr}(\Sigma_1) + \mu_1^\top\mu_1\right) \\ &= \frac{1}{2}\left(\log\prod_i \sigma_i^2 - n + \sum_i \sigma_i^2 + \sum_i \mu_i^2\right) \\ &= \frac{1}{2}\left(\sum \log\sigma_i^2 - n + \sum_i \sigma_i^2 + \sum_i \mu_i^2\right), \end{aligned} \tag{A-7}$$

where $|\cdot|$ indicates a determinant, and $\mu_i$ and $\sigma_i$ are the $i^{\text{th}}$ elements of $\mu_1$ and $\text{diag}(\Sigma_1)$ respectively.

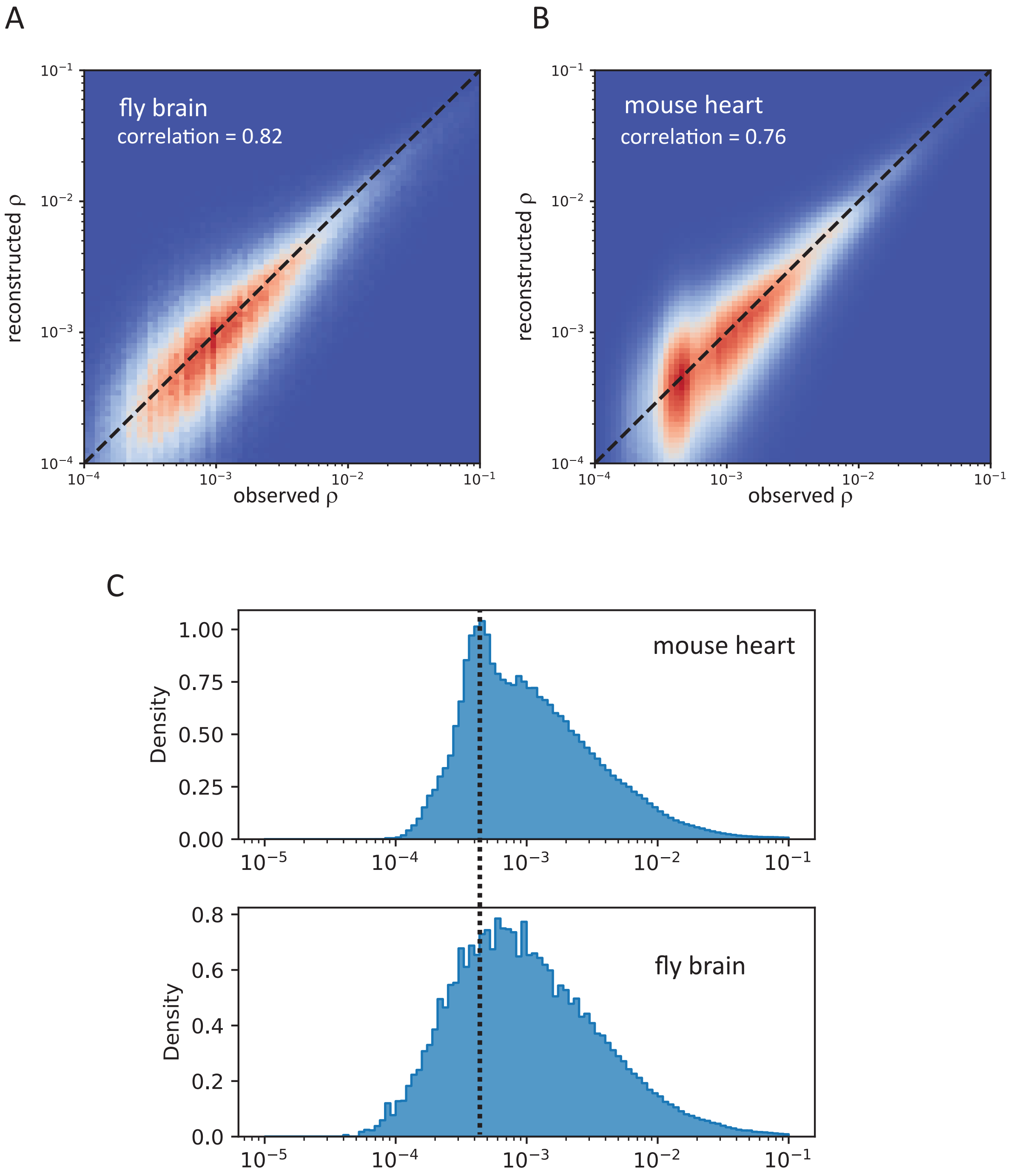


Figure S1: Relationship between the observed gene fraction and the reconstructed gene fraction for the clock neuron (A) and the mouse heart (B) dataset and their respective density (C).

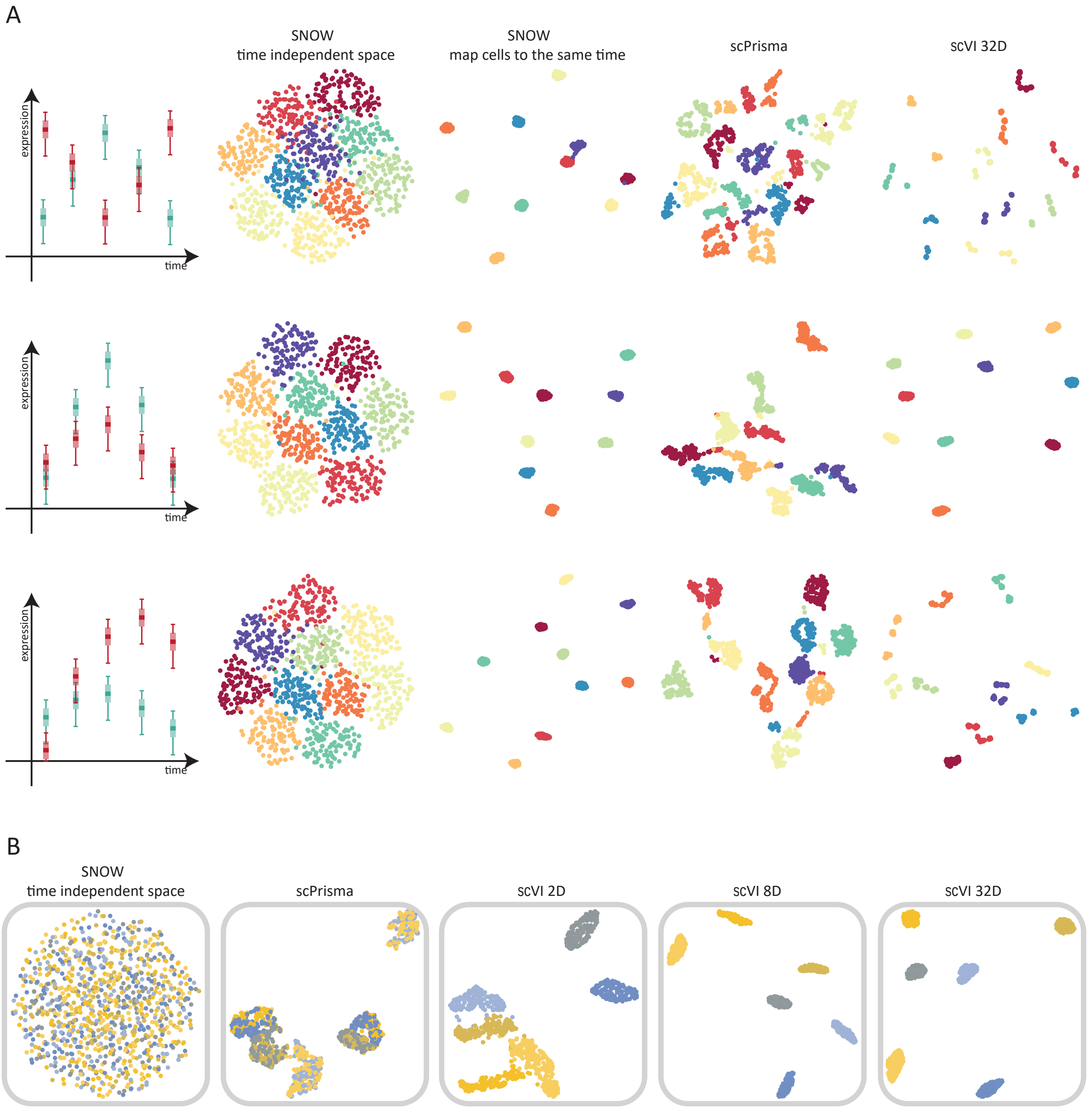


Figure S2: A: Low-dimensional projections of latent spaces generated by SNOW, scPrisma and scVI of the toy data. The toy data consists of 50 rhythmic genes and 100 flat genes. The rhythmic genes have a cell type–specific phase, cell type–specific amplitude, or a cell type–specific phase *and* amplitude in the first, second and third row respectively. Points are colored by cell type. The low-dimensional embeddings can be obtained in SNOW in two ways: either a UMAP plot of the latent space directly, or a UMAP plot of all cells projected (using the decoder) to a common time. The latter produces well–separated cell type clusters in all cases. B: Performance of SNOW on a toy dataset that contains only one cell type (and hence one cluster), colored by sample time. Low dimensional projections as generated using SNOW, scPrisma and scVI. For scVI we tested its performance using a 2D (scVI 2D), 8D (scVI 8D) and 32D (scVI 32D) latent space.

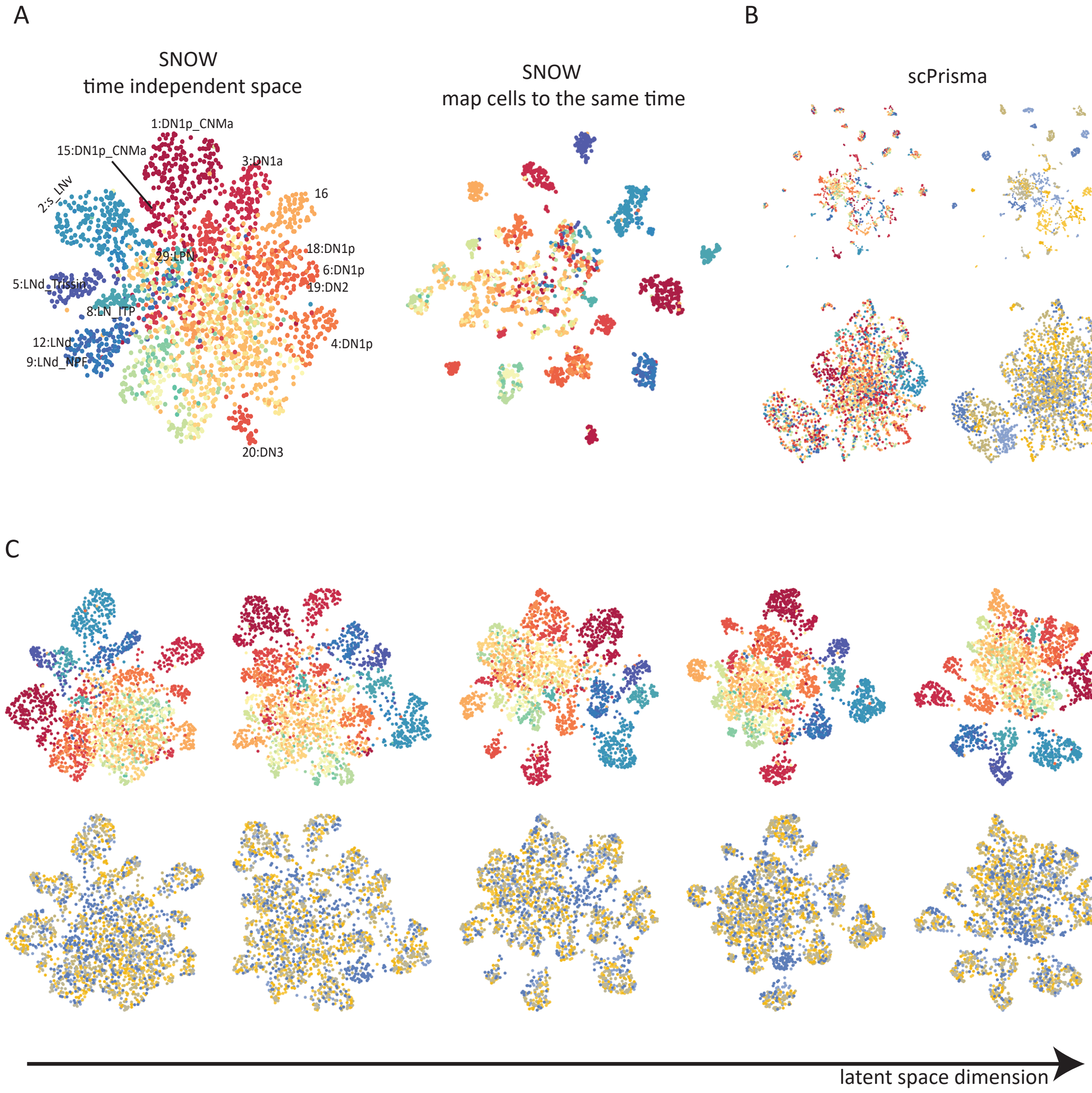


Figure S3: A: UMAP plots of the drosophila clock neuron data by using the SNOW time–independent space (left) or by mapping all the cells to the same time (right). B: scPrisma generated embedding of the drosophila clock neuron data by using log normalization (top row) and $Z$–scoring (bottom row). Colors on the left and right column denote cell type and time, respectively. C: scVI generated embedding of the drosophila clock neuron dataset using a 8D, 16D, 32D, 64D, or 128D latent space. Colors on the top and bootom denote cell type and time, respectively.

A

B

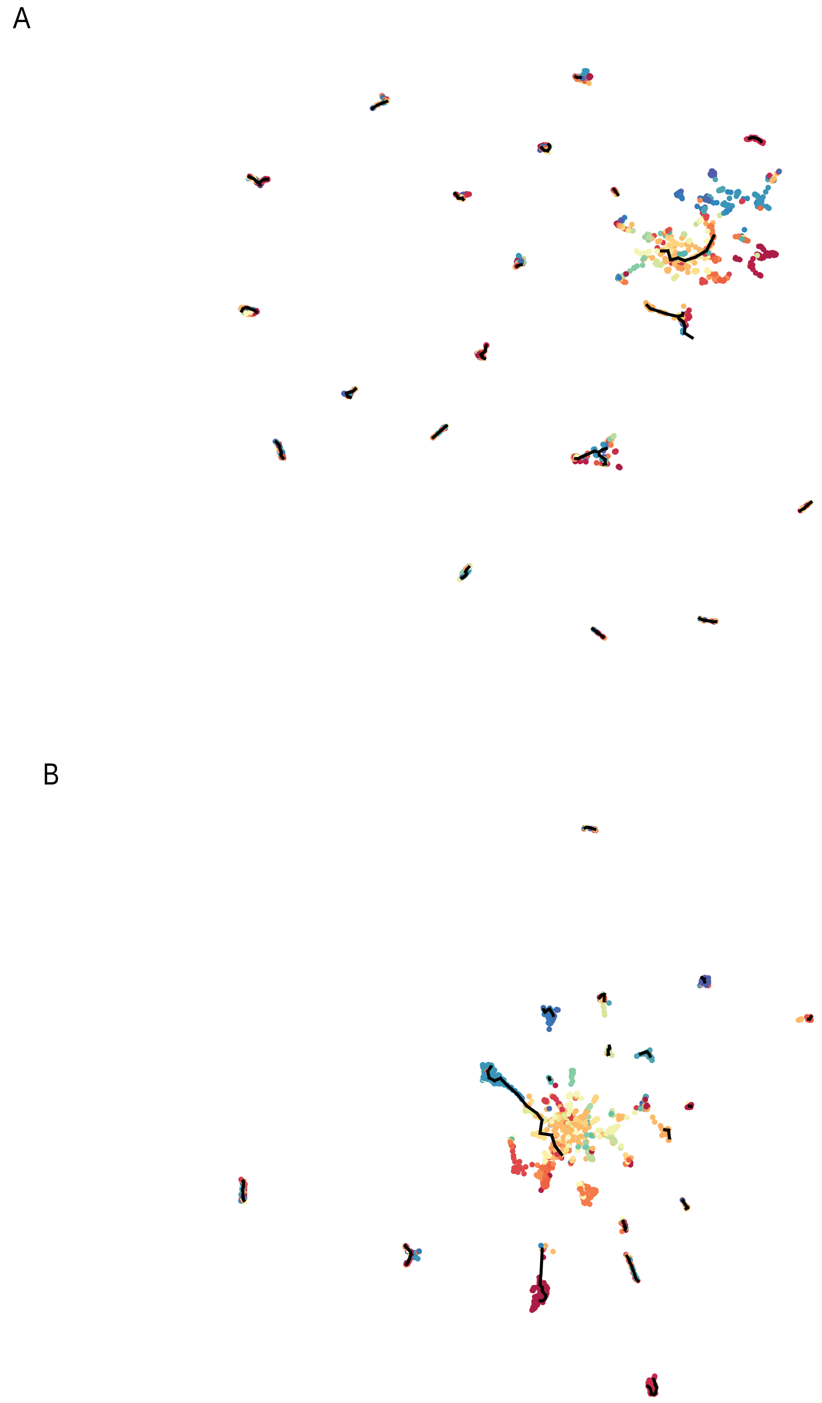

Figure S4: Cellular trajectories inferred from the drosophila clock neuron data using Monocle on the raw data (A) or batch normalized data (B).

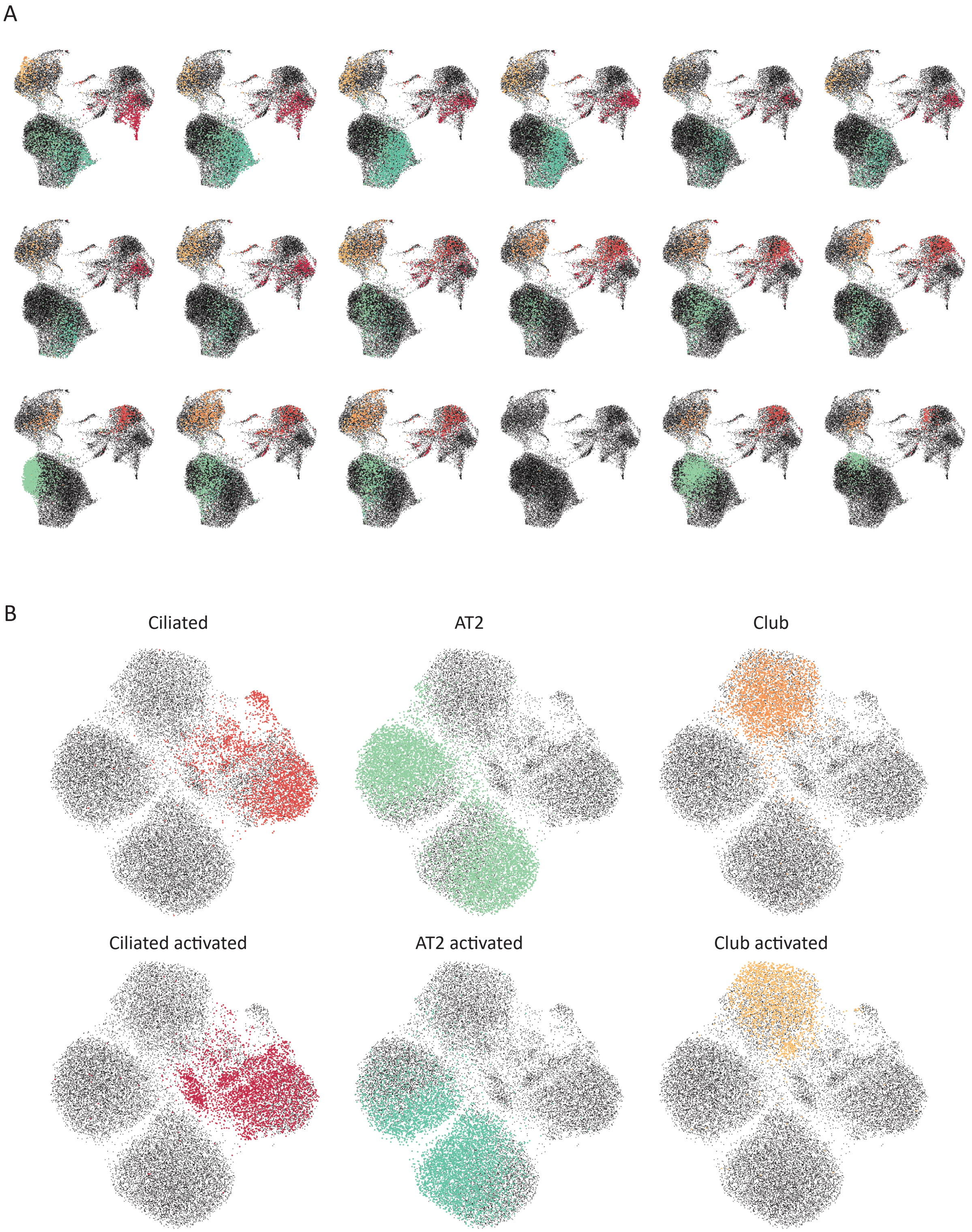


Figure S5: A: UMAP plots of lung regeneration data. Cells are collected every day for two weeks (days 1–14) and on days 21, 28, 36 and 54. Each panel shows all cells, with cells sampled on that particular day highlighted in color bu cell type, with other day's cells in black. Colors denote cell types. B: scVI–generated embedding of the lung regeneration dataset; for SNOW, see Figure S6.

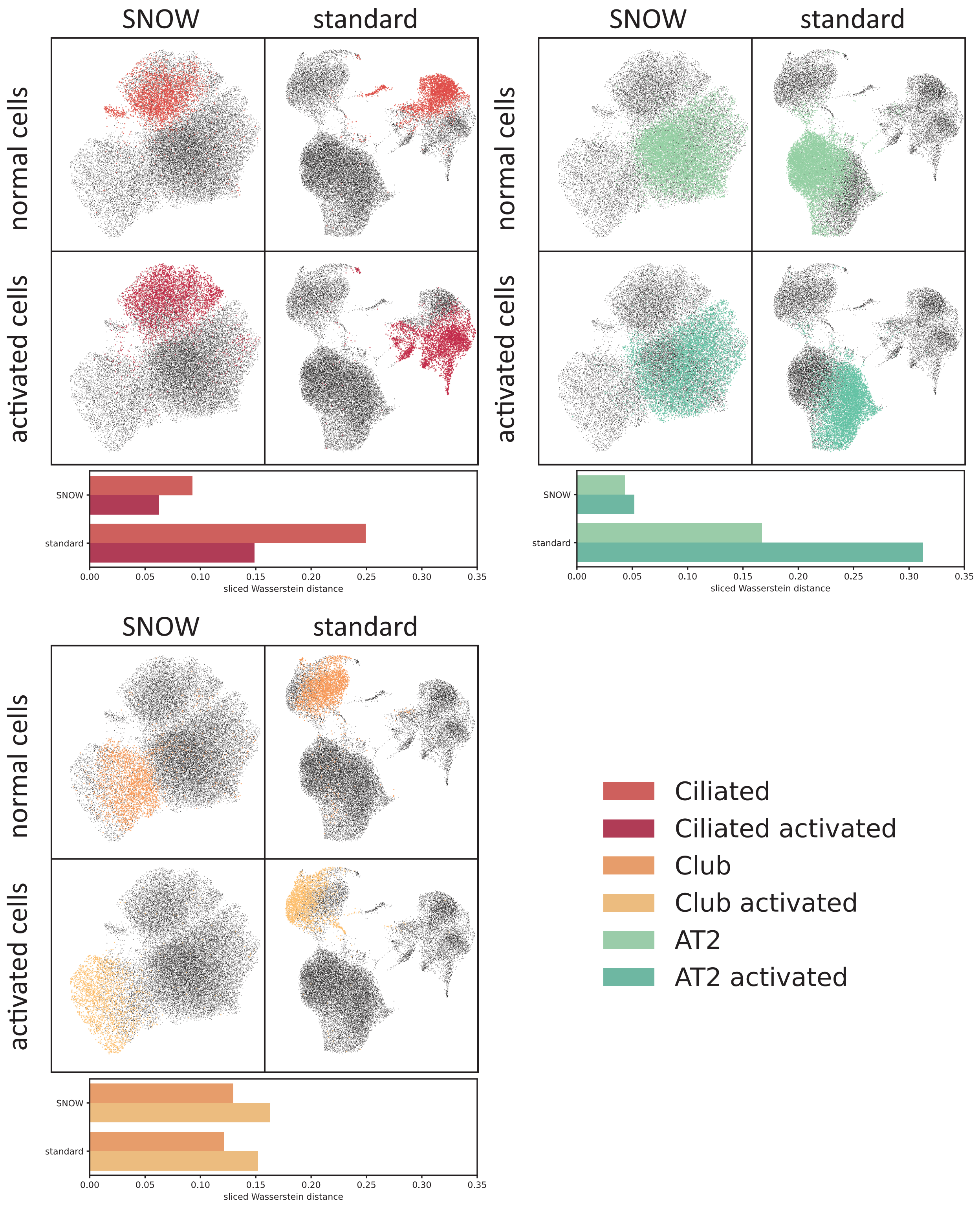


Figure S6: Comparison between the UMAP embedding produced with the raw data and with SNOW using the lung regeneration data. The bar chart below each panel shows the distance between each subtype and the main cell type it belongs to (for example, the distance between AT2 (subtype) and the combination of AT2 and AT2 activated). AT2: alveolar type 2 cells.

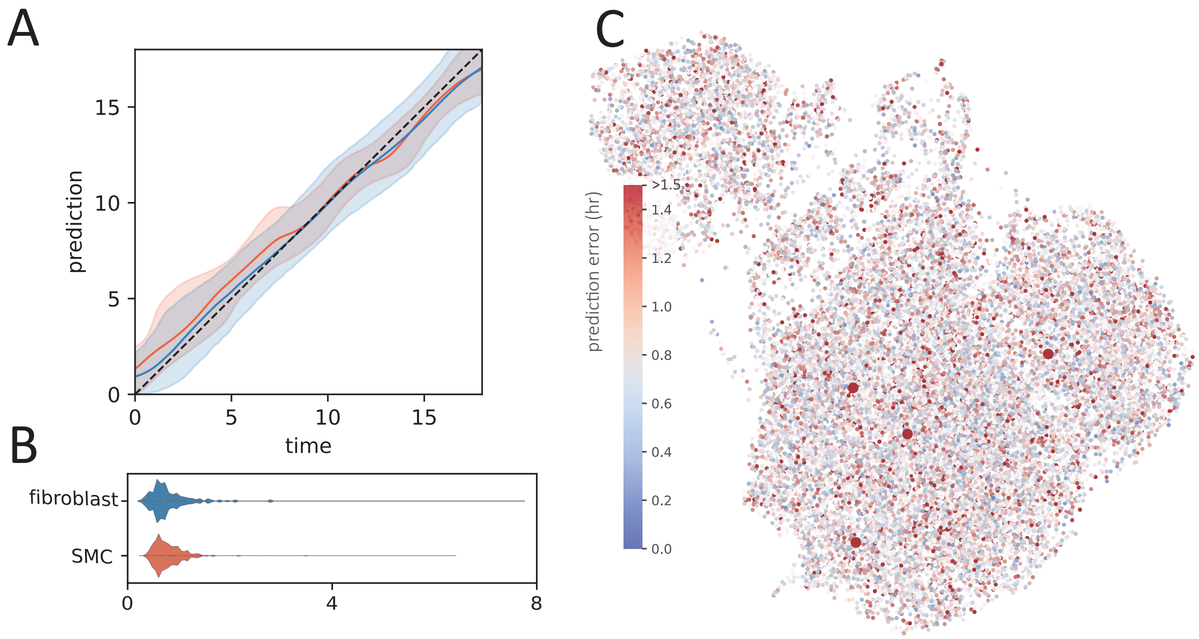


Figure S7: A: Relationship between the time inferred by the SNOW encoder ("prediction") vs the known time ("time") for SNOW–generated observations of mouse aorta cells at different times (blue: fibroblasts; orange: SMC). The shaded region indicates 95% confidence interval. B: Violin plots showing the mean absolute time prediction error of the fibroblast (blue) and SMC (orange) cluster. C: Mean absolute error overlayed on the UMAP projection of the mouse aorta data. Larger points indicate a mean absolute error greater than 5.

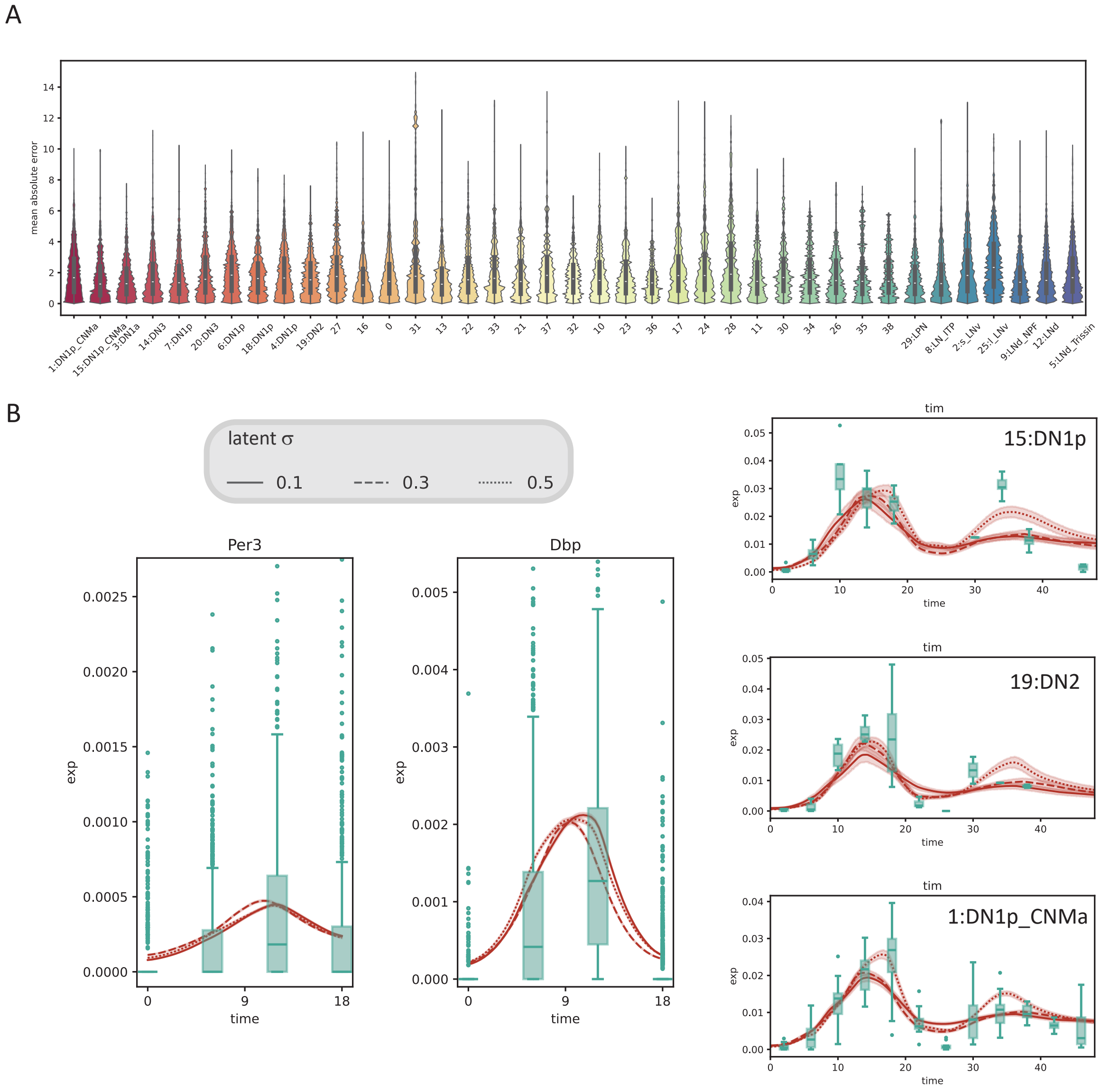


Figure S8: A: Mean absolute error of predicting time in the 38 neuron clusters from the clock neuron dataset. B: Impact of fixed latent space standard deviation on time series generation for the mouse heart (left) and the clock neuron dataset (right). Red traces indicate the mean of the generated data, with 95%CIs shaded. Green boxplots show the observed experimental data.

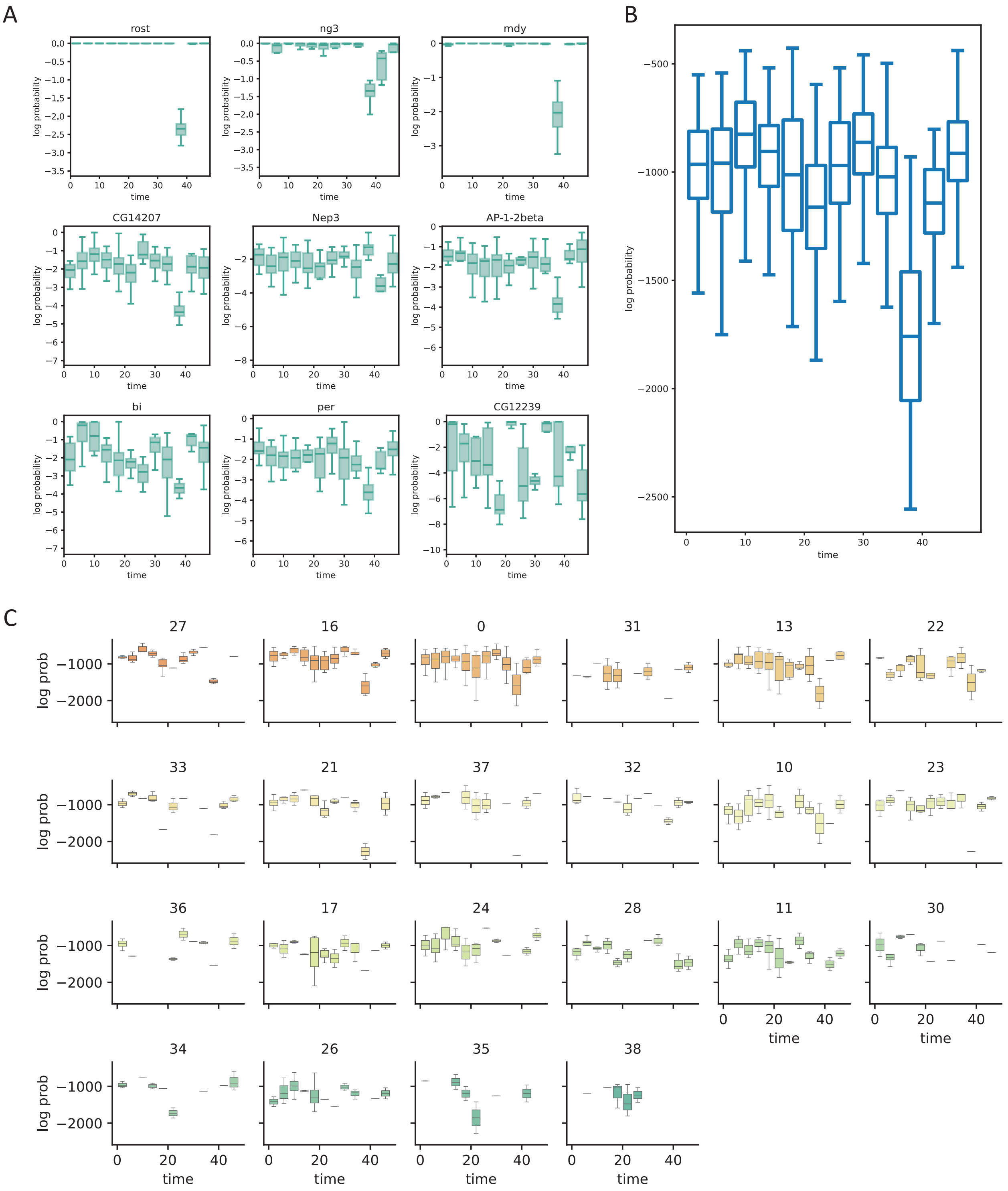


Figure S9: A: Box plot showing the log probability of observing each gene. B: Box plot showing the log probability of observing an entire cell. C: Box plot showing the log probability of observing each cell for the unnamed clusters.

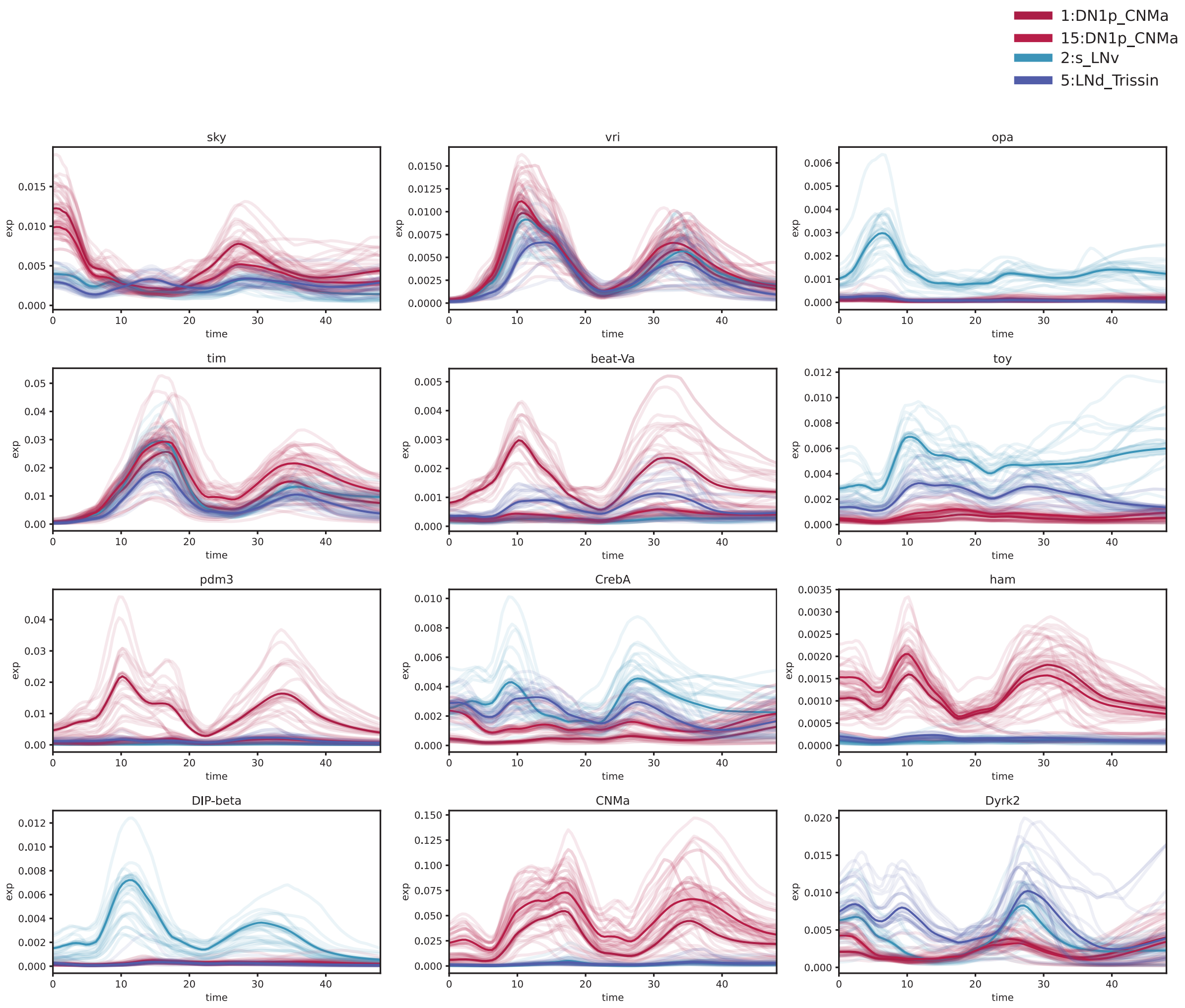


Figure S10: Examples of cell type–dependent gene expression dynamics. Colors denote different cell types.

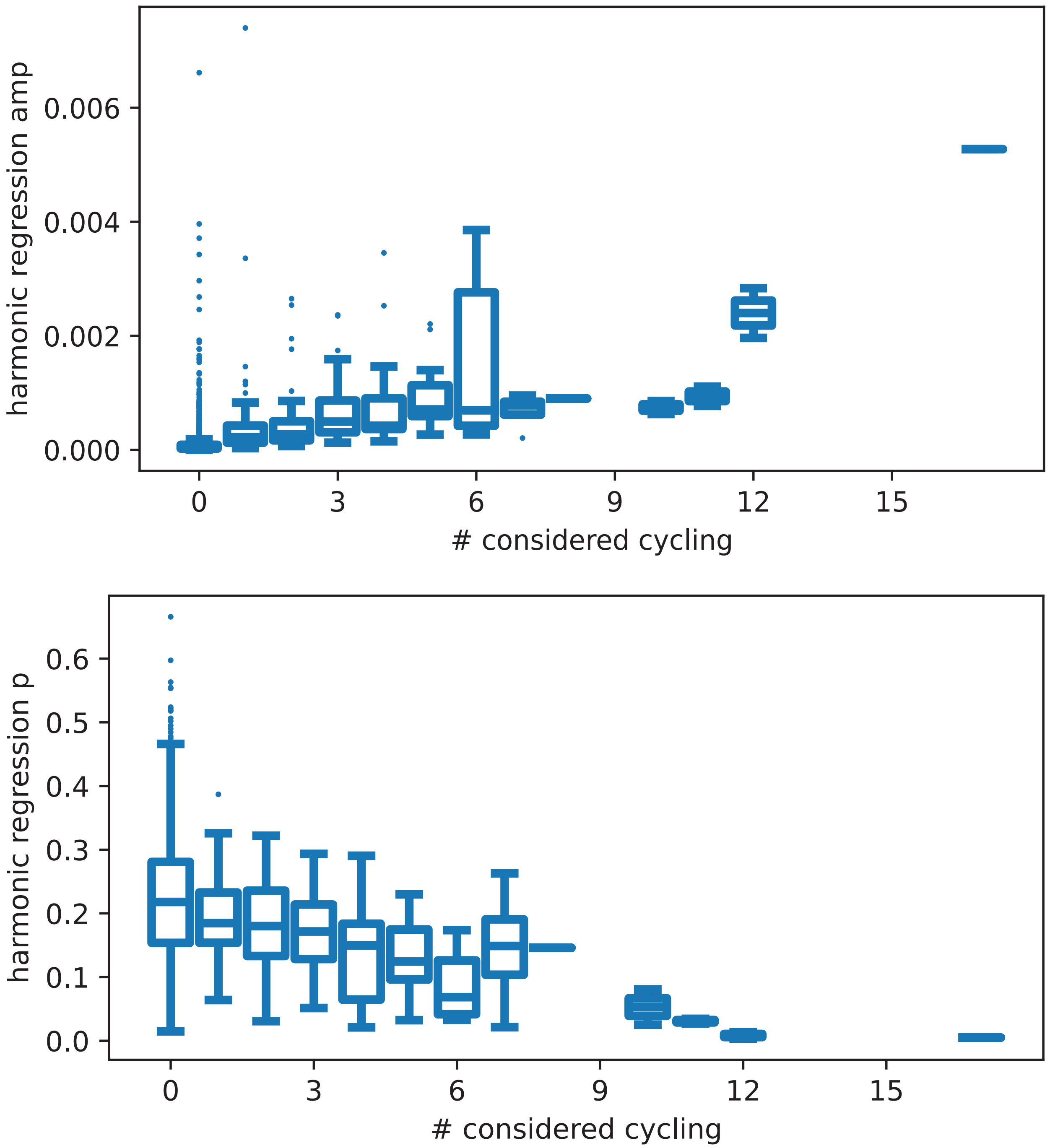


Figure S11: Top: Boxplots showing the relationship between the SNOW–estimated amplitude and the number of times a gene is reported to be cycling in [10]. Bottom: Boxplots showing the relationship between the cycling $p$-value from the SNOW–generated data and the number of times a gene is reported to be cycling in [10].

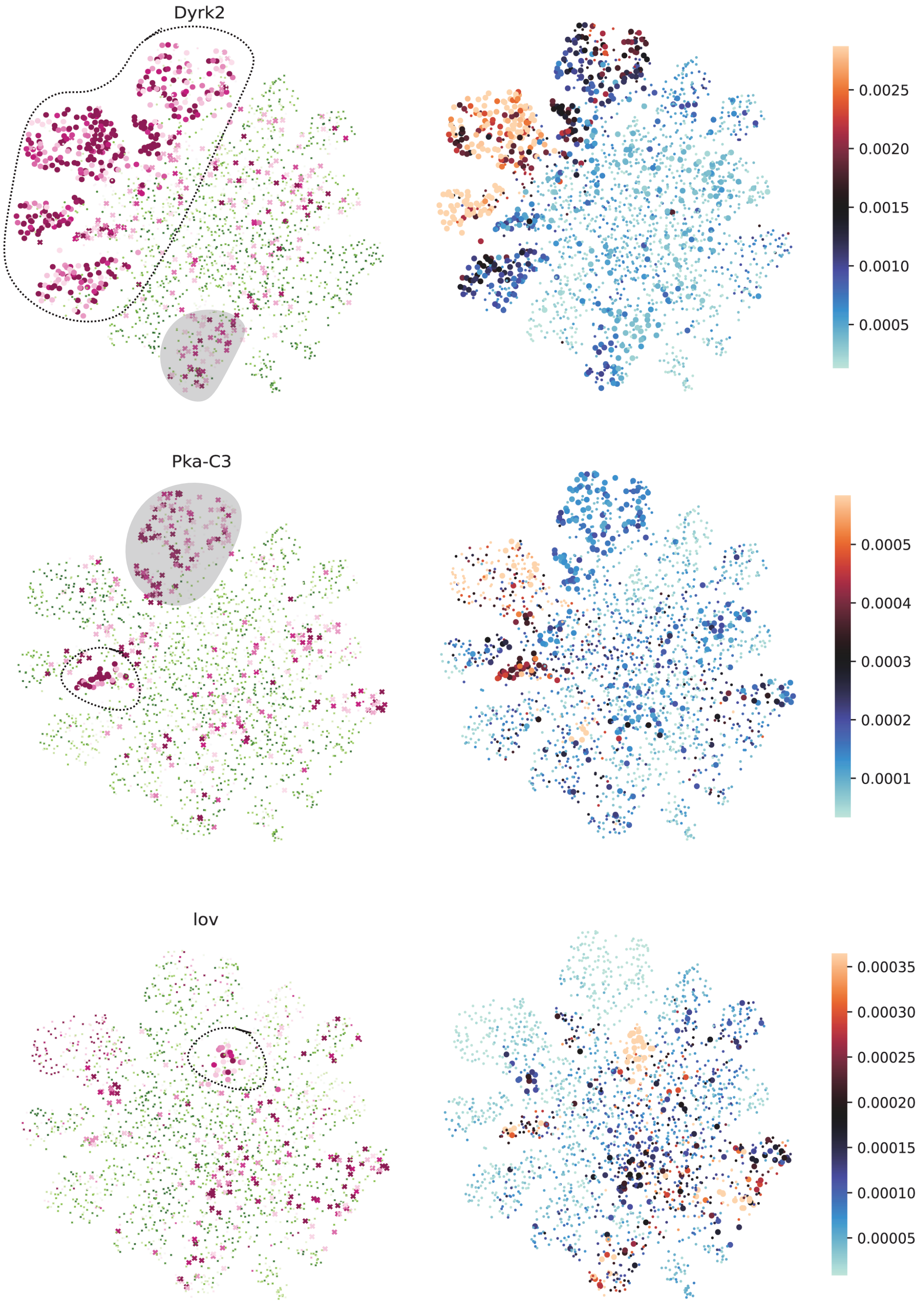


Figure S12: SNOW single–cell cycling analysis of genes reported as cycling in [10]. The left column shows estimated harmonic regression $p$-values and the right column shows estimated amplitudes. The circled region indicates agreement between our analysis and that of [10]; and the shaded region indcates disagreement. Cells with $p$-value greater than 0.001 or amplitude smaller than 0.0001 were made small for better visualization. Circular points indicate that this gene is reported to be cycling in the cell type this cell belongs to in [10], whereas crosses indicate that it is not considered cycling in [10].

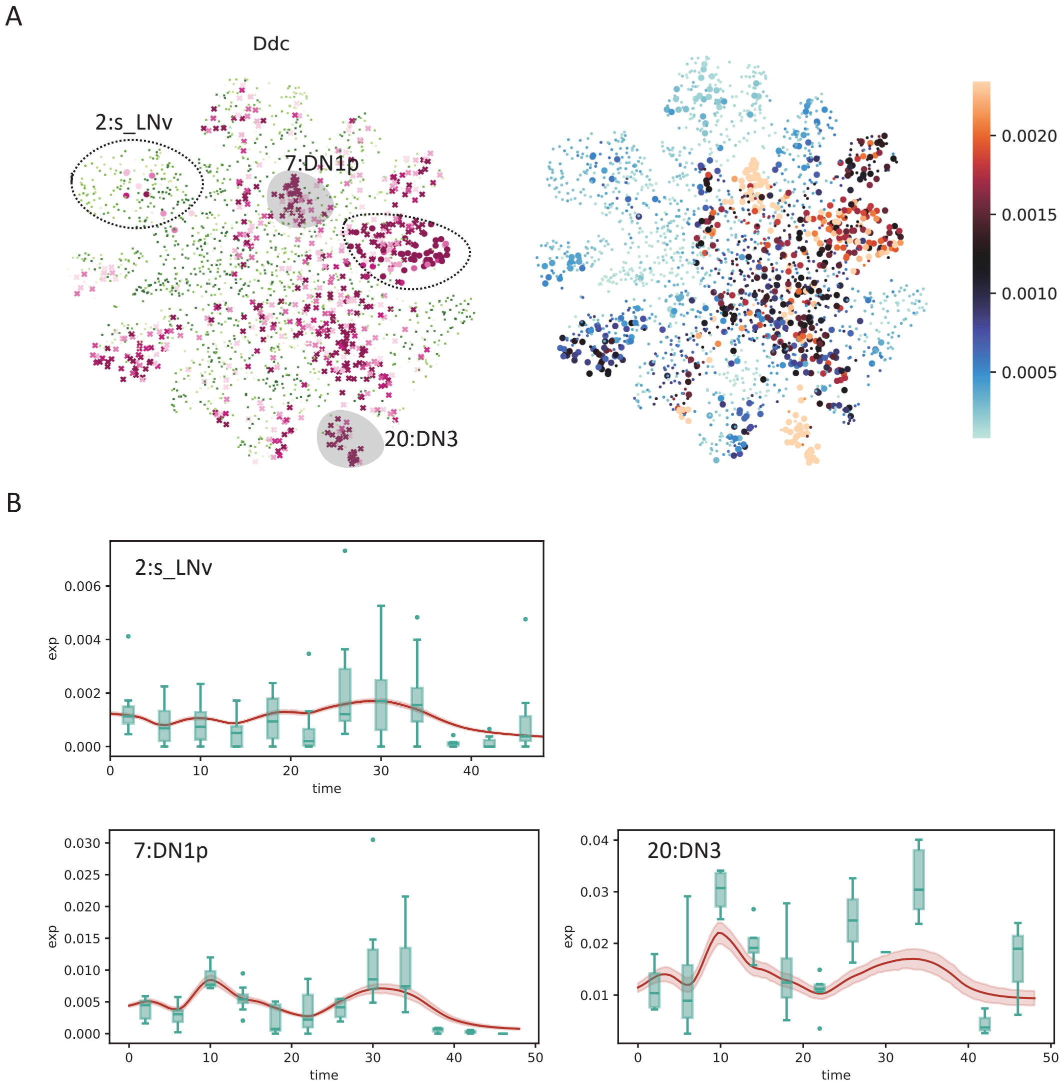


Figure S13: A: $p$-value and amplitude of *Ddc* overlayed on top of the UMAP projection of the clock neuron data. Colouring and points are as given in Figure S12. B: SNOW generated time series (average across cells shown as a red line, with shaded 95%CI) and experimental observations (green box plots) of cells within the circled and shaded regions of (A).
